# Mechanical fracturing of the extracellular matrix patterns the vertebrate heart

**DOI:** 10.1101/2025.03.07.641942

**Authors:** Christopher Chan Jin Jie, Daniel Santos-Oliván, Marie-Christine Ramel, Juliana Sánchez-Posada, Toby G R Andrews, Priscilla Paizakis, Emily S Noël, Alejandro Torres-Sánchez, Rashmi Priya

**Affiliations:** The Francis Crick Institute, 1 Midland Road, London, NW1 1AT, United Kingdom; European Molecular Biology Laboratory, EMBL Barcelona, C/ Dr. Aiguader 88, 08003 Barcelona, Spain; Barcelona Collaboratorium for Modelling and Predictive Biology, C/ Wellington 30, 08005 Barcelona, Spain; School of Biosciences and Bateson Centre, University of Sheffield, Western Bank, Sheffield, United Kingdom

**Author notes:** these authors contributed equally (Marie-Christine Ramel, Juliana Sánchez-Posada).

## Abstract

Pattern formation is fundamental to embryonic morphogenesis. In the zebrafish heart, spatially confined single-cell delamination in the ventricle outer curvature initiates trabeculation, a conserved morphogenetic process critical for heart function and embryonic life. Yet, what confines delamination in the ventricle outer curvature remains ill-understood. Contrary to the prevailing notion of patterning through biochemical signals, we now show that mechanical fracturing of the cardiac extracellular matrix (cECM) patterns delamination in the outer curvature. cECM fractures emerge preferentially in the outer curvature, cells delaminate into these fractures and experimental blocking of fractures blocks delamination. These fractures display characteristic signature of mechanical defects and myocardial tissue contractility is sufficient to fracture the cECM, independent of molecular signals, enzymatic activity, or delamination events. Notably, the anisotropic geometry of myocardial tissue generates higher mechanical strain in the outer curvature, thereby locally patterning cECM fractures and delamination. Consequently, cECM fractures evolve in response to dynamic changes in tissue geometry, and experimental manipulation of tissue geometry is sufficient to alter the fracture pattern. Together, our findings underscore mechanical fractures as a morphogenetic strategy, and more generally, corroborate the long-standing but understudied paradigm that tissue form-function can feed back to steer its own patterning.

## Main

During embryonic development, localized patterning of cellular processes in space and time generates intricate morphological structures to sculpt functional organs. In the last decades, developmental patterning has been primarily explained by genetic prepatterning (*1, 2*). However, tissue patterning is a multiscale emergent phenomenon guided by macroscopic mechanical constraints, including tissue geometry and extracellular environment (*3-7*). Further, many developing organs like heart, gut and lungs undergo significant mechanical deformations as part of their physiological function, which in turn can influence patterning across scales, from individual cell fate to branching morphogenesis (*8-10*). Yet, mechanistically, how these global mechanical cues orchestrate localized cellular behaviours to steer robust tissue patterning remains elusive.

During the first 72 hours of development, the zebrafish heart undergoes remarkable changes in its geometry while experiencing dynamic mechanical forces generated by its own rhythmic contractions (*11-15*). These macroscale transformations are coupled with spatially confined single-cell delamination events that initiate trabeculation—a morphogenetic process critical for heart contraction and blood flow (*13, 16-18*). This, combined with its tractability and accessibility, makes the developing zebrafish heart an excellent system for investigating how the geometry and function of a developing organ feeds back to its patterning.

The primitive zebrafish heart begins as a linear tube that loops into an S-shape to form two chambers, atrium and ventricle (*11, 12, 14, 15*). The ventricle myocardial tissue expands anisotropically, acquiring a bean shape with a distinct bulge, broadly referred to as the outer curvature (OC, **Fig. 1A**). Subsequently, to support the increasing functional demands of the heart, trabeculation initiates specifically in the outer curvature—a feature conserved across vertebrate hearts, including humans (*13, 18-24*). During trabeculation, the myocardial tissue transitions from a simple monolayer into an intricate three-dimensional (3D) meshwork, consisting of two distinct cell types – outer compact layer (CL) cells enveloping an inner layer of contractile trabecular cells (TL) (*4, 13, 16, 17*) (**Fig. 1A**). Trabeculae are multicellular myocardial ridges, which increase the muscle mass of the embryonic heart to ensure efficient blood flow, heart contraction and facilitate nutrient/oxygen uptake (*13, 16, 18, 19, 25*). Anomalous trabecular morphogenesis leads to embryonic lethality and cardiomyopathies in humans (*13, 19, 25, 26*), thus emphasising the fundamental importance of studying trabecular morphogenesis for health and disease.

**Fig. 1.**
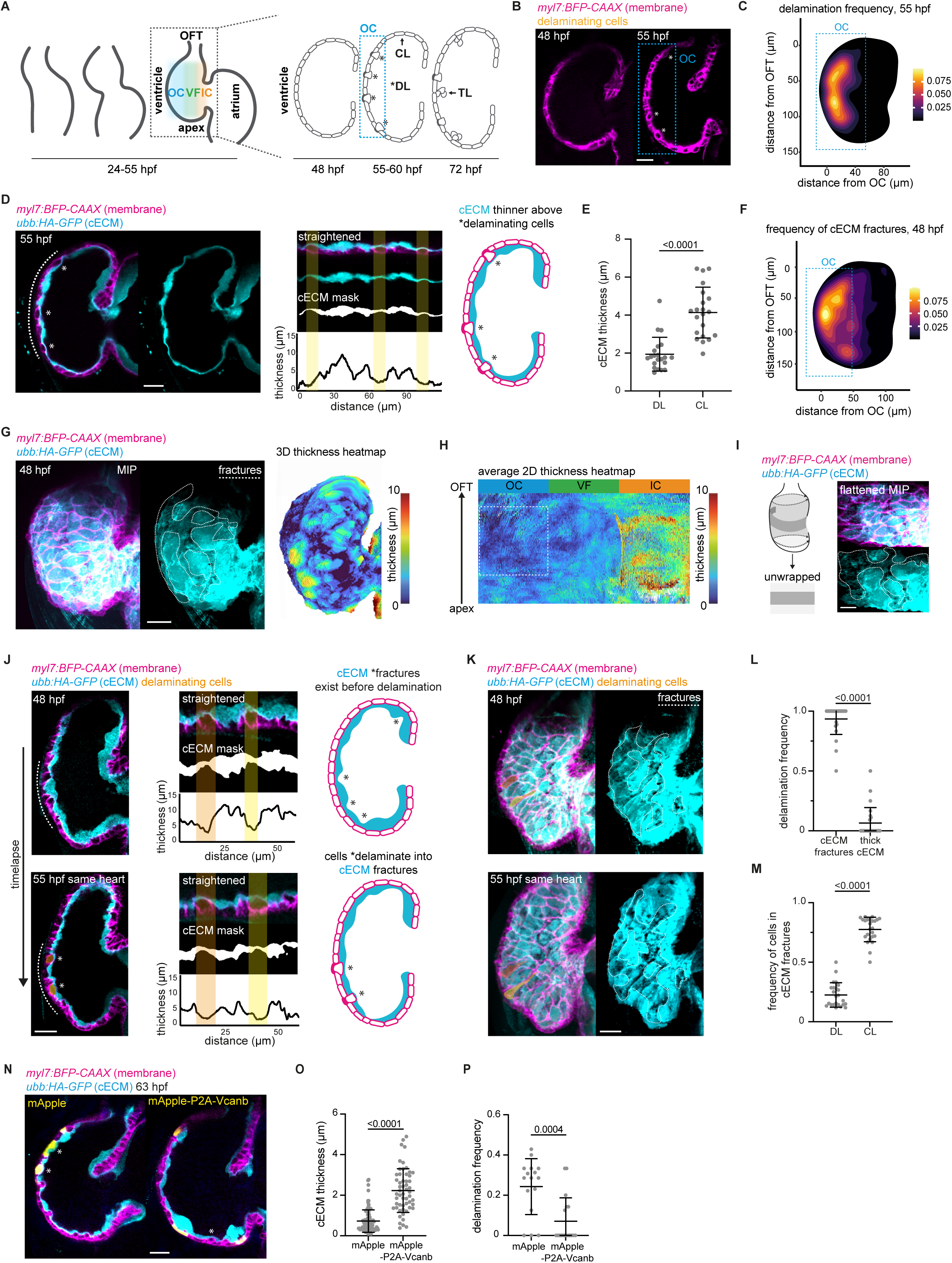
cECM fractures spatially restrict delamination in the ventricle outer curvature. **(A)** Schematic showing changes in heart geometry coincide with localized delamination (DL) in the outer curvature (OC) of the ventricle, which initiates trabeculation (TL). CL, compact layer; VF, ventral face; IC, inner curvature; OFT, outflow tract. **(B-C)** Midsagittal section of a 48 hpf ventricle showing monolayer myocardial tissue, n = 43/43 embryos, and a 55 hpf ventricle showing onset of delamination in the outer curvature (OC, cyan box); delaminating cells marked by asterisks, n = 37/37 embryos **(B)**, frequency heatmap of delamination at 55 hpf; cyan box, OC, n = 29 embryos **(C)**. *myl7:BFP-CAAX* marks cell membrane of cardiomyocytes. **(D-E)** Midsagittal sections of a 55 hpf ventricle and the corresponding 2D cECM thickness map from the marked area (white dotted line), shaded yellow lines indicate overlap of thin cECM (fractures) and delamination, schematic showing that cECM is thinner above delaminating (DL) cells **(D),** quantification of cECM thickness above delaminating (DL) versus compact layer (CL) cells; n = 21 embryos **(E)**. *ubb:HA-GFP* labels cECM. **(F)** Frequency heatmap of cECM fractures, defined as less than 2 µm, in a 48 hpf ventricle; cyan box, outer curvature (OC). n = 16 embryos. **(G-I)** Maximum intensity projection (MIP) of a 48 hpf ventricle, fractures highlighted using dotted line; corresponding 3D cECM thickness heatmap, n = 41/41 embryos **(G)**, and average 2D cECM thickness heatmap generated using morphoHeart, n = 8 embryos; OC, dotted line **(H)**, and corresponding flattened MIP showing cECM fractures are independent of the shapes and sizes of the overlying myocardial cells **(I)**. **(J-K)** Midsagittal sections from longitudinal imaging of the same ventricle at 48 hpf and 55 hpf, delaminating cells are pseudo coloured, representative of n = 49/53 delaminating cells from 20 embryos, 2D cECM thickness map from the marked areas (white dotted line) with yellow and orange shaded lines indicating overlap of cECM fractures and delamination, and schematic showing cells delaminating into cECM fractures **(J)**, corresponding maximum intensity projection (MIP) with delaminating cells pseudo coloured showing cells delaminate into cECM fractures (white dotted regions) **(K)**. **(L)** Frequency of cells delaminating into cECM fractures versus thick cECM regions, n = 24 embryos. **(M)** Frequency of delaminating (DL) versus non-delaminating or compact layer (CL) cells in cECM fractures, n = 24 embryos. **(N-P)** Midsagittal section of a 63 hpf ventricle overexpressing mApple and Vcanb **(N),** quantification of cECM thickness underlying mApple expressing cells (n = 94 cells) and Vcanb expressing cells (n = 55 cells) **(O)** and delamination frequency of mApple (n = 16 embryos) or Vcanb expressing cells (n = 18 embryos) **(P)**. Asterisks mark delaminating cells. All data are mean ± s.d. P values result from unpaired two-tailed Students’ t-tests. Scale bars, 20µm.

In zebrafish, trabeculation begins around 55 hours post-fertilization (hpf) and is triggered by single cell delamination driven by local heterogeneity in actomyosin tension (*17*). A subset of cells with higher actomyosin tension constrict their apical domain and stochastically delaminate towards the lumen to seed the inner trabecular layer (*16, 17*) (DL, **Fig. 1A-B**). Numerous reports suggest that delamination starts in the outer curvature of the ventricle (*16, 17, 20, 27, 28*). However, this phenomenon has only been qualitatively studied due to the intricate 3D structure of the myocardial wall, which complicates the analysis of single-cell behaviour. Accordingly, we developed a morphometric pipeline to quantitatively analyse delamination in 3D (**Fig. S1A**) and established that delaminating cells are indeed confined to the ventricle outer curvature at the onset of trabeculation (55 hpf) (**Fig. 1B-C**). Yet, why delamination starts in the outer curvature remains ill-understood.

A long-held theory is that the preferential activation of a key upstream signal, Nrg2a-Erbb2, specifies trabecular cells in the outer curvature of vertebrate hearts (*13, 16-18, 20-22, 25, 27-30*). To test if Nrg2a-Erbb2 signalling is indeed enriched in the outer curvature, we first quantified *erbb2* and *nrg2a* expression using multiplexed hybridization chain reaction (HCR) (*31*). Our analysis suggests that neither transcript is enriched in the outer curvature (**Fig. S1B-D**). We then examined Erbb2 activity by quantifying the activity of its canonical downstream effector ERK (*21, 32*) through live imaging of an ERK-KTR biosensor (*33*), and found that ERK activity is also not enriched in the outer curvature (**Fig. S1E-G**). Overall, these results indicate that Nrg2a-Erbb2 signalling is not enriched in the outer curvature, thus ruling out its role in patterning delamination specifically in the outer curvature.

To address what patterns delamination, we turned our focus to the extracellular matrix (ECM), as ECM remodelling has been shown to guide trabecular morphogenesis in vertebrates (*18, 26, 34, 35*), and is an important regulator of tissue patterning (*36-38*). We began by analysing the basement membrane components and asked whether there is a spatiotemporal correlation between delamination and basement membrane remodelling. A careful analysis of basement membrane components Laminin and Fibronectin before (48 hpf) and at the onset of delamination (55 hpf) revealed no spatial correlation between delamination and localization of these BM components (**Fig. S2A-E**). At 48 hpf, Fibronectin is enriched in the valve endocardium (**Fig. S2A**) while Laminin localizes to the basal side of the myocardium (**Fig. S2B**), as previously reported (*39, 40*). Moreover, at 55 hpf, we did not observe any changes in Fibronectin localization (**Fig. S2C**), and Laminin localization does not spatially correlate with delamination events (**Fig. S2D-E**), indicating that basement membrane remodelling is not required to induce or pattern delamination.

We then focussed on the cardiac jelly, a layer of cardiac extracellular matrix (cECM), which is important for various aspects of vertebrate heart morphogenesis, including trabeculation (*18, 26, 34, 35, 41-43*). cECM is primarily composed of hyaluronic acid and is sandwiched between the endocardium and myocardium (**Fig. S2F**) (*34, 44, 45*). We started by visualizing spatiotemporal dynamics of cECM in live hearts using a well-characterized biosensor line which labels hyaluronic acid (*34, 41, 44, 45*). Strikingly, we observed a strong spatial correlation between cECM thinning and delamination events, as cECM is ∼ 50% thinner above delaminating cells (∼ 2 µm) (**Fig. 1D-E**). Additionally, cECM thickness is already heterogenous as early as 48 hpf (∼ 7 hrs before the onset of delamination), indicating that cECM thinning precedes delamination (**Fig. S2G**). We further confirmed these observations using an orthogonal approach by analysing the negative space between the myocardium and endocardium, as cECM is tightly sandwiched between these tissue layers. We find that cECM thickness is indeed heterogenous at both 48 and 55 hpf (**Fig. S2H-I**). Notably, our morphometric analysis revealed a conspicuous spatial correlation between cECM thinning and delamination in the outer curvature of the ventricle. These thin cECM regions are already biased in the outer curvature of the ventricle at 48 hpf, before the onset of delamination (**Fig. 1F**), and importantly, this strongly correlates with subsequent delamination in the outer curvature at 55 hpf (**Fig. 1C**).

Next, to quantitatively analyse cECM thickness, we utilized a 3D segmentation pipeline, morphoHeart (**Fig. S2J**) (*46*). Using this approach, we derived 3D thickness heatmaps of the cECM, enabling holistic analysis of cECM thickness in the context of heart geometry (**Fig. 1G-H**). These 3D thickness maps were then unrolled and superimposed to generate average 2D heatmaps to quantitatively represent average cECM thickness distribution across multiple embryos (**Fig. 1H, S2J**). We found that cECM displays heterogenous distribution at 48 hpf, further confirming that cECM thinning precedes delamination, and that thin cECM regions are prominently localized in the outer curvature (**Fig. 1G-H**). Notably, these thinner areas are independent of the shapes and sizes of the overlying myocardial cells, and appear as fractures in the cECM, visible as irregular breaks running across the myocardial tissue with length-scales greater than a single cell (**Fig. 1I, S2K-L, Supplementary Video 1**). Together, we conclude that these thin cECM regions, which we hereafter refer to as cECM fractures, are enriched in the outer curvature before the onset of delamination.

These observations prompted us to investigate whether cells delaminate into cECM fractures. Using longitudinal live imaging, we tracked cECM fractures and delamination events in the same heart and found that some cECM fractures at 48 hpf were indeed subsequently occupied by delaminating cells at 55 hpf (**Fig. 1J-K, Supplementary Video 2**). Next, using mosaic labelling of cells with mCardinal to track single cell behaviour in live hearts, we find that myocardial cells send protrusions into cECM fractures (**Fig. S2M**), indicating that cells can sense local cECM fractures. Strikingly, although majority (∼ 90%) of cells delaminate into cECM fractures (**Fig. 1L**), not all cECM fractures are occupied by delaminating cells (**Fig. 1M**), suggesting that cECM fractures are permissive, rather than selective for delamination.

To further strengthen the notion that cECM fractures facilitate delamination, we developed a genetic tool to locally thicken the cECM and subsequently analysed its effect on delamination. We expressed Versican b mosaically specifically in cardiomyocytes. Versican b interacts with hyaluronan and has glycosaminoglycan chains which allows it to sequester water and ions (*43, 47, 48*). We reasoned that mosaic expression of Versican b may lead to local swelling of the hyaluronan matrix, thereby thickening the hyaluronan-rich cECM. Indeed, cells expressing Versican b have a significantly thicker layer of underlying cECM, and correspondingly, their delamination is abrogated (**Fig. 1N-P**). Notably, this phenotype is maintained at 72 hpf (**Fig. S2N-P**), suggesting that delamination is not delayed, but indeed abrogated. Together, we conclude that cECM fractures facilitate delamination in the ventricle outer curvature.

Given its importance in patterning delamination, we then asked what leads to cECM fracturing. In mice, Nrg-Erbb2 and Notch signalling regulates cECM remodelling during trabeculation (*18*). Accordingly, we analysed cECM distribution in Erbb2 inhibited (**Fig. 2A-B**), *nrg2a* mutant (**Fig. S3A-B**), and Notch inhibited hearts (**Fig. S3D-E**) and found that cECM fractures still form, thus indicating that neither Nrg-Erbb2 nor Notch signalling are required to generate cECM fractures in zebrafish. We then asked whether delaminating cells could actively remodel their local cECM. To test this hypothesis, we abrogated delamination by blocking the upstream signal Nrg-Erbb2 (*16, 17, 27, 28*). A careful analysis of Erbb2 inhibited and *nrg2a* mutant hearts at 55 hpf revealed that while they lacked delamination, cECM fractures still form (**Fig. S3C,F**). We then induced ectopic delamination in hearts by increasing actomyosin contractility of single cells using constitutively active myosin light chain 9 expression, as done previously (*17*). We find that ectopic delamination does not locally redistribute cECM (**Fig. 2C-D**), further confirming that cECM fractures are not caused by delamination. Together, these results suggest that cECM fracturing is not induced by delamination, Nrg-Erbb2 signalling or Notch signalling.

**Fig. 2.**
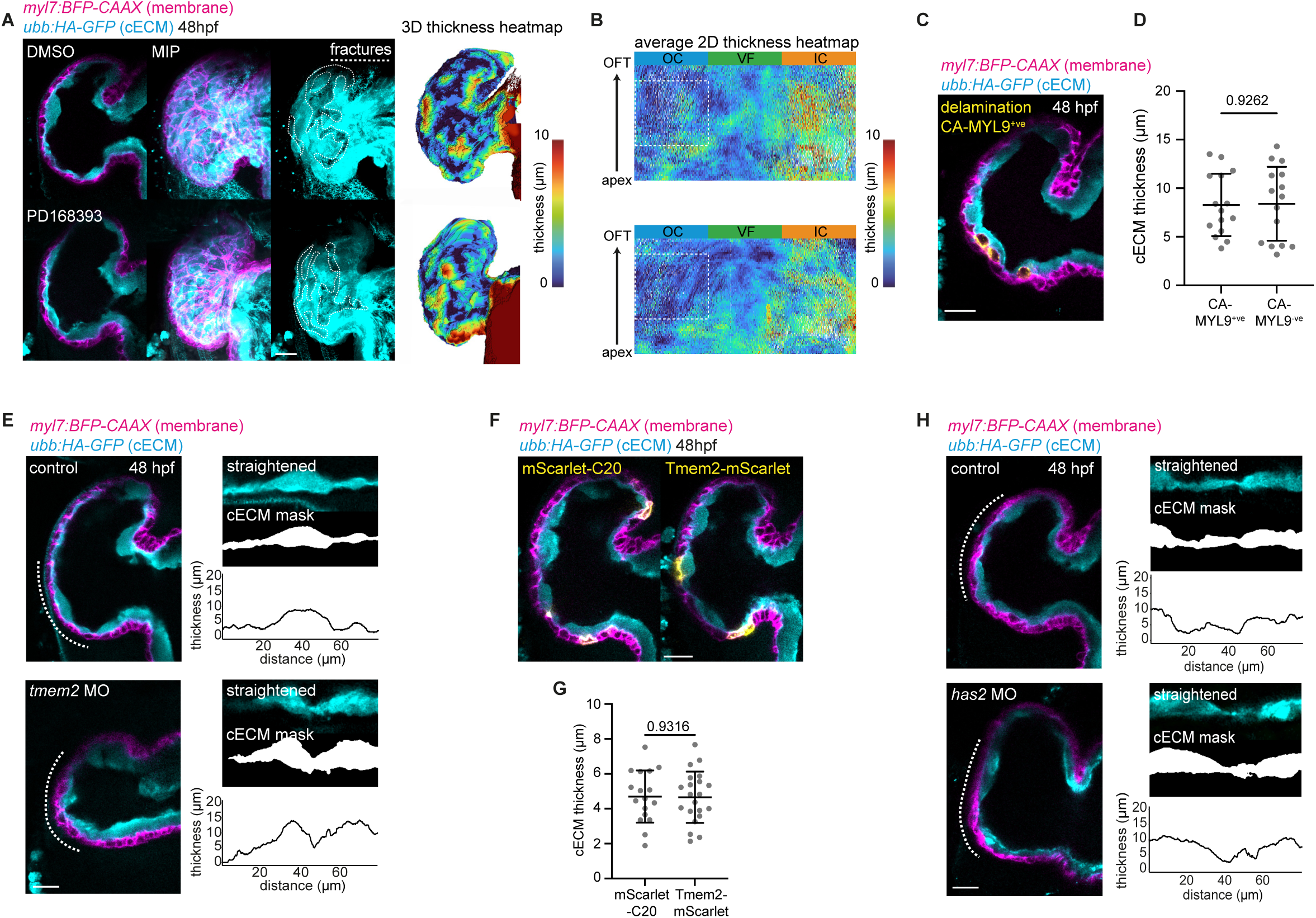
cECM fractures emerge independently of Nrg2a-Erbb2 signalling, delamination and enzymatic activity. **(A-B)** 48 hpf ventricles from DMSO (control) and Erbb2 inhibited (PD168393) embryos, n = 17/17 embryos each, corresponding 3D cECM thickness heatmap **(A)**, and average 2D cECM thickness heatmaps generated with morphoHeart, n = 7 embryos each **(B)**. *myl7:BFP-CAAX* marks the cell membrane of cardiomyocytes, *ubb:HA-GFP* labels cECM. **(C-D)** 48 hpf ventricle from embryos injected with constitutively active MYL9 (CA-MYL9) to induce ectopic delamination **(C)**, and cECM thickness above CA-MYL9^+ve^ and CA-MYL9^-ve^ neighbouring cells, n = 15 embryos **(D)**. **(E)** cECM distribution in 48 hpf control (n = 15 embryos) and *tmem2* morphant (MO) ventricles (n = 18 embryos) and corresponding 2D cECM thickness map from the marked areas (dotted lines). **(F-G)** 48 hpf ventricles expressing mScarlet-C20 (n = 17 embryos) or Tmem2-mScarlet (n = 21 embryos) **(F)**, and quantified cECM thickness **(G)**, showing no changes in cECM thickness. **(H)** cECM distribution in 48 hpf control (n = 13 embryos) and *has2* morphant (MO) ventricles (n = 14 embryos) and corresponding 2D thickness map from the marked areas (white dotted lines). All data are mean ± s.d. P values result from unpaired two-tailed Students’ t-tests. Scale bars, 20µm.

We then asked whether local enzymatic synthesis or degradation of cECM is responsible for generating cECM fractures. We first inhibited matrix metalloproteases (MMPs) using a broad range inhibitor GM6001 and found that cECM fractures persist (**Fig. S4A**), confirming that MMPs are not required to generate cECM fractures. As the cECM is primarily composed of hyaluronan (*42, 44, 45, 49, 50*) we reasoned that hyaluronidases that are specifically expressed in the ventricle could locally degrade the cECM to fracture it. Utilizing in-situ hybridization to do a broad screen of candidate genes, we found that *cemip2* (*tmem2*) and *hyal2b* appear to be enriched in the ventricle at 48 hpf (**Fig. S4B**). To further quantitatively analyse their expression at the cellular level, we performed hybridisation chain reactions and found that neither *tmem2* (**Fig. S4C-E**) nor *hyal2b* transcripts (**Fig. S4F-H**) are enriched in the outer curvature or localize to cECM fractures. As Tmem2 has been shown to be the main hyaluronidase responsible for degradation (*45, 51, 52*), we further examined its role in generating cECM fractures using gain-and loss-of-function approaches. Using a morpholino against *tmem2* to reduce its protein levels, we find that heart looping is affected as previously reported (*53*) but cECM fractures are still present (**Fig. 2E**). We then performed mosaic overexpression of Tmem2 specifically in cardiomyocytes and found that this was not sufficient to fracture the cECM (**Fig. 2F-G)**. Next, we examined the role of hyaluronan synthases (*has2*), the predominant hyaluronan synthase expressed in the heart (*41*) and found that its expression is restricted to the valves (**Fig. S4I**). Further, although *has2* morphants have looping defects and thinner cECM in the valves as previously reported (*54*) (**Fig. 2H**), cECM fractures still form, indicating that local synthesis also does not contribute towards heterogenous cECM organisation. Together, these results indicate that local enzymatic synthesis or degradation is not required to generate cECM fractures, which is also in line with previous findings that the total cECM volume does not decrease during the early stages of heart development (*35, 46*). Collectively, we conclude that cECM fractures form independent of Nrg2a-Erbb2 signalling, Notch signalling, delamination, or enzymatic degradation and synthesis.

We then reasoned that to understand how cECM gets fractured, we needed a deeper understanding of its spatiotemporal dynamics. However, *in situ* live imaging of the zebrafish ventricle at early stages (< 2 dpf) at cellular resolution still remains difficult due to the tissue depth and orientation of the heart which obstructs analysis of early ventricle morphogenetic events (*41*). To this end, we developed a body-explant method, where we surgically removed the anterior region of embryos and allowed them to recover in supplemented L15 culture medium for one hour (**Fig. S5A, Supplementary Video 3**). We find that body explants reestablish systemic circulation and maintain morphologically as well as functionally normal hearts (**Fig. S5B, Supplementary Video 3**).

Utilizing this method, we were able to image entire 24 hpf beating hearts, *in situ* and at cellular resolution (**Fig. 3A, Supplementary Video 4**). Interestingly, we found that the cECM appears relatively homogenous at 24 hpf, at the onset of heart contraction (**Fig. 3A, Supplementary Video 5**). However, at 30 hpf (∼ 24 hours before the onset of delamination) cECM fractures start forming, as areas of thin and thick cECM were clearly visible (**Fig. 3B, Supplementary Video 5**). These results suggest that cECM fractures emerge concomitantly as the heart develops and reinforces our earlier observation that delamination does not induce cECM fracturing. Notably, at 30 hpf, fractures extend linearly across the myocardial tissue along the heart’s major axis (**Fig. S5C**) resembling mechanical fractures. In a pressurized cylindrical sheet undergoing cyclic expansion—such as the 30 hpf heart—strain develops perpendicular to the cylinder’s major axis which will cause fractures to appear along the major axis, consistent with our *in vivo* observations (*55*). These findings suggest that a tissue-scale mechanical process may be driving cECM fracturing. We therefore examined tissue-scale mechanical parameters that change between 24-34 hpf, when the cECM fractures emerge.

**Fig. 3.**
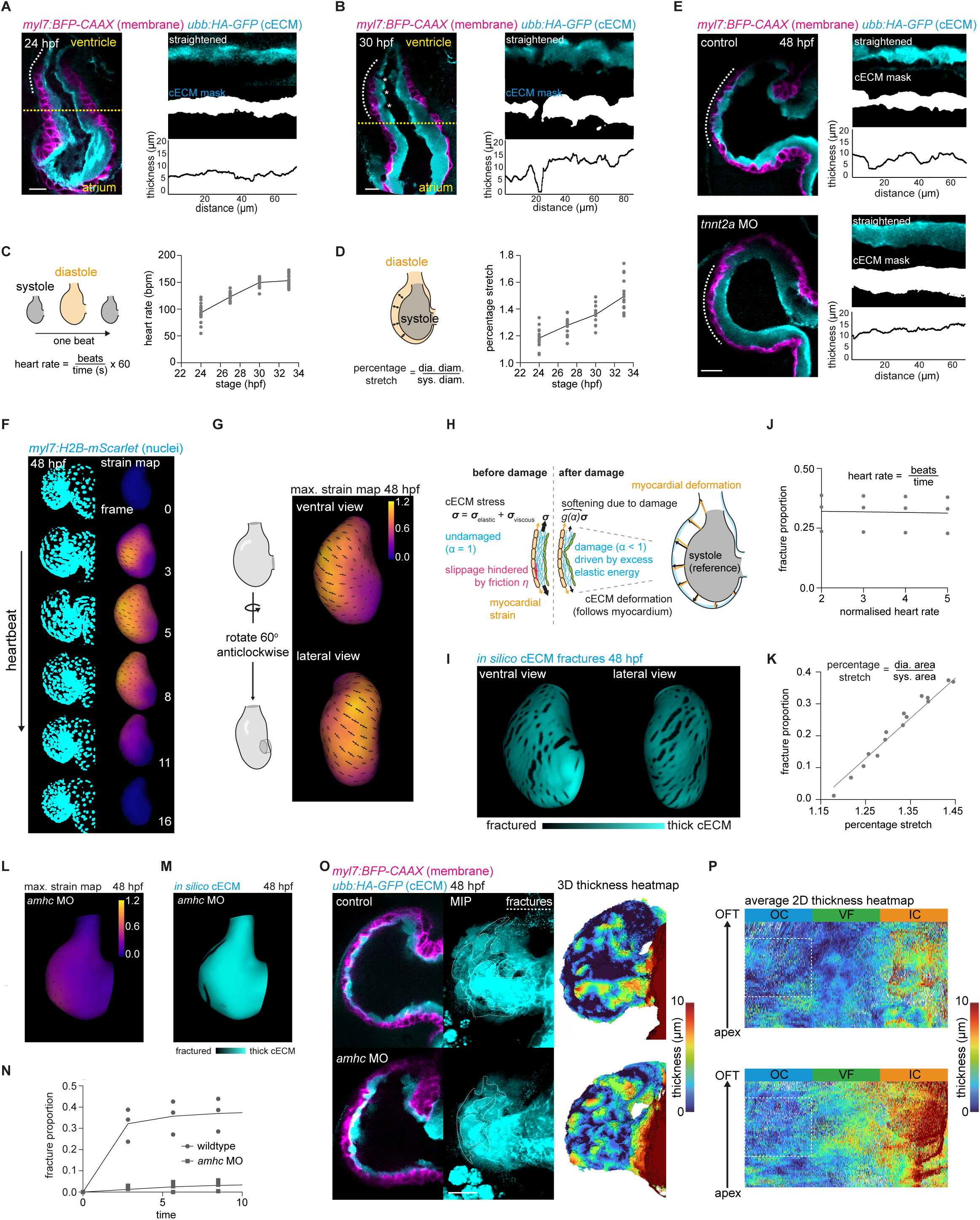
cECM fracturing is mechanically driven by myocardial tissue stretching. **(A-B)** Distribution of cECM in a 24 hpf ventricle showing homogenous thickness and corresponding 2D cECM thickness map from the marked areas (dotted lines), n = 26/28 embryos **(A)**, distribution of cECM in a 30 hpf ventricle showing fractures emerging (asterisks), and corresponding 2D cECM thickness map from the marked areas (dotted lines), n = 15/17 embryos **(B).** *myl7:BFP-CAAX* marks cell membrane of cardiomyocytes, *ubb:HA-GFP* labels cECM. **(C-D)** Explanatory schematics and quantification of heart rate **(C)** and extent of myocardial tissue stretch (percentage stretch) **(D)** between 24 hpf to 34 hpf, n = 15 embryos/stage. **(E)** cECM distribution in a 48 hpf control, n = 20 embryos, and *tnnt2a* morphant (MO) ventricle, n = 21 embryos, and corresponding 2D cECM thickness map from the marked areas (dotted lines). **(F)** Myocardial tissue strain maps calculated across a heart cycle in a 48 hpf ventricle, *myl7:H2B-mScarlet* marks the nuclei of cardiomyocytes. **(G)** Ventral and lateral views of maximum strain maps for a 48 hpf ventricle. Arrows indicate the directions of maximum and minimum strain, and their size represents the magnitude of the strain. **(H-I)** Biophysical model of the damage and fracturing of the cECM, for more details, please see modelling section **(H),** ventral and lateral view of a 48 hpf heart showing cECM fracture patterns obtained from simulations **(I)**. **(J-K)** Simulation predictions showing effect of heart rate **(J)** and percentage stretch **(K)** on cECM fracturing predicting that proportion of cECM fractures increases with percentage stretch. **(L-N)** Maximum strain maps **(L)** and *in silico* cECM fractures pattern **(M)** of a 48 hpf *amhc* morphant (MO) ventricle showing reduced strain and fractures, respectively. Graph showing reduced cECM fracture proportion in *amhc* MO ventricle compared to wild type ventricle **(N)**. Arrows in **(L)** indicate the directions of maximum and minimum strain, and their size represents the magnitude of the strain. **(O-P)** Midsagittal view and maximum intensity projection (MIP) showing cECM distribution in a 48 hpf control (n = 19 embryos) and *amhc* morphant (MO) ventricle (n = 23 embryos), corresponding 3D cECM thickness heatmap **(O),** and average 2D cECM thickness heatmap derived using morphoHeart, n = 6 embryos each **(P)**. All line graphs (except for **3K**) are plotted following the mean of the data. Scale bars, 20µm.

We observed a significant increase in both heart rate and the extent the myocardial tissue stretches during heart beats between 24-34 hpf (**Fig. 3C,D**). We thus hypothesized that mechanical forces arising from the increasing heart rate and tissue stretch might fracture the cECM. To test this hypothesis, we blocked heart contraction genetically using a *tnnt2a* morpholino (*16, 17, 20, 27*) (**Fig. 3E**) and chemically using a calcium channel blocker, Nifedipine (*56*) (**Fig. S5D**). Strikingly, blocking myocardial tissue contraction completely abrogates cECM fracturing, leading to a homogenous cECM thickness using both approaches. To confirm that cECM fractures are specifically triggered by ventricular contraction, we analysed *half-hearted* mutants in which the atrium contracts, but ventricular contraction is abrogated (*57*). Consistent with our previous findings, we find that the cECM in *half-hearted* ventricles is not fractured, thus underscoring the role of ventricular contraction in inducing cECM fractures (**Fig. S5E**). Since manipulating heart contractility can influence blood flow patterns and, in turn, shear stress (*58*), we asked whether shear stress could also drive cECM fracturing. Accordingly, we analysed *gata1* morphants which lack circulating blood cells, and as a result have significantly reduced shear stress (*59*) and found that cECM fractures still form in the outer curvature (**Fig. S5F-G)**. Together, these results argue that cECM fracturing is a mechanical process driven mainly by myocardial tissue contractility.

We then asked whether this mechanical remodelling of cECM might represent a general phenomenon observed in other vertebrate hearts. To test this hypothesis, we analysed the cECM dynamics of chick embryos as they are tractable for live imaging (*60*). At HH10, when the heart tube initially forms, the heart is not yet beating, and the cECM appears uniform (**Fig. S6A**), similar to the 24 hpf zebrafish heart. Interestingly, at the onset of heart contractions around HH12, the cECM begins to redistribute and fracture-like patterns emerge (**Fig. S6A**). Notably, when heart contractions are inhibited using Nifedipine, this redistribution is absent, and the cECM remains thick and uniform (**Fig. S6B**), similar to what we observe upon stopping contractions in zebrafish hearts. Given that heterogeneities in cECM thickness is observed across vertebrate hearts (*18, 26, 35, 42, 44, 45, 49, 50*), it is tempting to speculate that cECM mechanical fracturing might be a common mechanism driving vertebrate heart morphogenesis

Next, to gain a deeper insight into the dynamics of cECM fracturing and how heart contractility regulates this process, we developed a quantitative biophysical model. We extracted deformation dynamics of the beating heart by tracking myocardial nuclei (**Fig. 3F, S7A, Supplementary Video 6**) and used this to reconstruct myocardial strain maps, which represent the extent and orientation of local tissue stretch (**Fig. 3F-G, S7B, Supplementary Video 7**). We then imposed this myocardial deformation onto a simulated cECM, modelling the cECM as an isotropic viscoelastic sheet with frictional interaction with the myocardium which undergoes persistent damage due to cyclic loading (**Fig. 3H**, see supplementary modelling section). To corroborate that the zebrafish cECM behaves as a viscoelastic solid as has been previously shown for the chick cECM (*42, 61*), we stopped hearts for extended periods of time and observed that heterogeneities in cECM thickness do not dissipate (**Fig. S7C**). Moreover, the cECM is also capable of retaining its shape in decellularized zebrafish hearts (**Fig. S7D**) similar to chick hearts (*43*).

Interestingly, applying cyclic myocardial deformation to the cECM in simulations induces fractures that first nucleate as regions of slightly damaged cECM, which then deepen and become more pronounced over time (**Fig. 3I, Supplementary Video 8**), consistent with the behaviour of fractures in embryos (**Fig. S7E**). Notably, our model predicted that cECM fracturing is primarily driven by the extent of myocardial stretch, while rate of contraction (heart rate) has limited effect (**Fig. 3J-K**). To test this prediction, we first used orthogonal pharmacological approaches to manipulate heart contractility, which affects both heart rate and myocardial tissue stretch. IBMX and Isoprenaline augments contractility while Nifedipine and Lidocaine abrogates contractility (*24, 56*). Consistent with our modelling predictions, quantitative analysis of cECM distribution in 48 hpf embryos revealed more cECM fractures when contractility is augmented and less cECM fractures when contractility is abrogated (**Fig. S7F-G**).

Next, to decouple the effects of heart rate from myocardial stretch, we used *amhc* morphants. In the absence of *amhc*, the atrium does not beat while the rate of ventricle contractions is not compromised (*62*) (**Fig. S7H**). Since the atrium is unable to pump blood into the ventricle, we reasoned that the *amhc* morphant ventricle myocardium will undergo less stretching, thereby allowing us to decouple ventricular contraction rate from its stretch. Quantification of myocardium deformation dynamics in *amhc* morphant ventricles reveal that indeed the ventricle undergoes less stretching (**Fig. 3L, S7I, Supplementary Video 9).** Correspondingly, our model predicted that cECM fracturing will be reduced in the *amhc* morphant ventricles (**Fig. 3M-N, Supplementary Video 10**). Consistent with our modelling predictions, analysis of cECM distribution in *amhc* morphant embryos confirm that cECM fracturing is reduced, suggesting that the extent of myocardial stretch is primarily responsible for cECM fracturing (**Fig. 3O-P**).

Since cECM fractures permit delamination (**Fig. 1J-M**), we then asked if manipulating fractures will affect delamination efficiency. To test this, we pharmacologically manipulated heart contractility and thereby cECM fracturing as described previously and analysed delamination at 55 hpf. Abrogation of heart contractility by Nifedipine and Lidocaine leads to less cECM fractures and correspondingly the number of delaminating cells is significantly lower than the untreated hearts (**Fig. S7J-K**). Interestingly, although treatment with IBMX and Isoprenaline increases heart contraction and thereby cECM fracturing, these hearts do not have more delaminating cells (**Fig. S7J-K**). This suggests that ectopic induction of cECM fracturing does not lead to ectopic delamination, confirming our previous observations that cECM fractures are permissive rather than selective. Together, we conclude that cECM fracturing is driven by myocardial stretching deformations and is required for delamination.

Notably, in our model, cECM fractures are enriched in the outer curvature of the ventricle (**Fig. 3I, S8A**), consistent with our *in vivo* observations (**Fig. 1F**). A careful analysis of the myocardial tissue strain maps revealed that the outer curvature experiences the greatest deformation, resulting in the highest strain (**Fig. 3G**), thereby spatially constraining cECM fractures in the outer curvature (**Fig. 3I, S8A**). Given that the tissue geometry has been postulated to pattern mechanical and chemical signals to guide morphogenesis (*4, 5, 63-69*), we hypothesized that the maximum strain in the outer curvature arises from the anisotropic geometry of myocardial tissue. This implies that myocardial strain maps and cECM fractures should be patterned differently at earlier stages of heart development when geometry is simpler, and that these patterns should evolve as the heart undergoes dynamic geometric transformations between 30 and 42 hpf.

To test this hypothesis, we computed tissue strain maps from nuclear tracks of early beating hearts (**Fig. S8B-D**) and integrated these into our biophysical model. At 30 hpf, when the heart is relatively linear, the maximum strain localizes predominantly to the dorsal and ventral sides. As the ventricle undergoes geometric changes, maximum strain shifts accordingly and gets concentrated in the outer curvature (**Fig. 4A, Supplementary Video 11**). Additionally, the orientation of maximum strain evolves over time. At 30 hpf, maximum strain aligns perpendicular to the heart’s long axis and becomes increasingly isotropic at 42 hpf (**Fig. 4A**). Concomitantly, our model predicts that at 30 hpf cECM fractures form on the dorsal and ventral sides of the heart and are oriented along its major axis (**Fig. 4B, S8E**). As the heart changes its geometry, new cECM fractures emerge in the outer curvature, aligning perpendicular to the direction of maximum strain (**Fig. 4B, S8E, Supplementary Video 12)**. Consistent with these predictions, as myocardial geometry evolves, the localization of cECM fractures also shifts *in vivo* (**Fig. 4C-D**). In a 30 hpf heart, fractures are observed on the dorsal and ventral sides, mirroring strain maps and simulation predictions (**Fig. 4D**). As the heart undergoes looping and adopts an anisotropic bean shape, fractures become increasingly concentrated in the outer curvature of the ventricle (**Fig. 4D**). Furthermore, as predicted by simulations, fractures at 30 hpf align predominantly with the heart’s major axis but display a more complex orientation at later developmental stages (**Fig. 4C**).

**Fig. 4.**
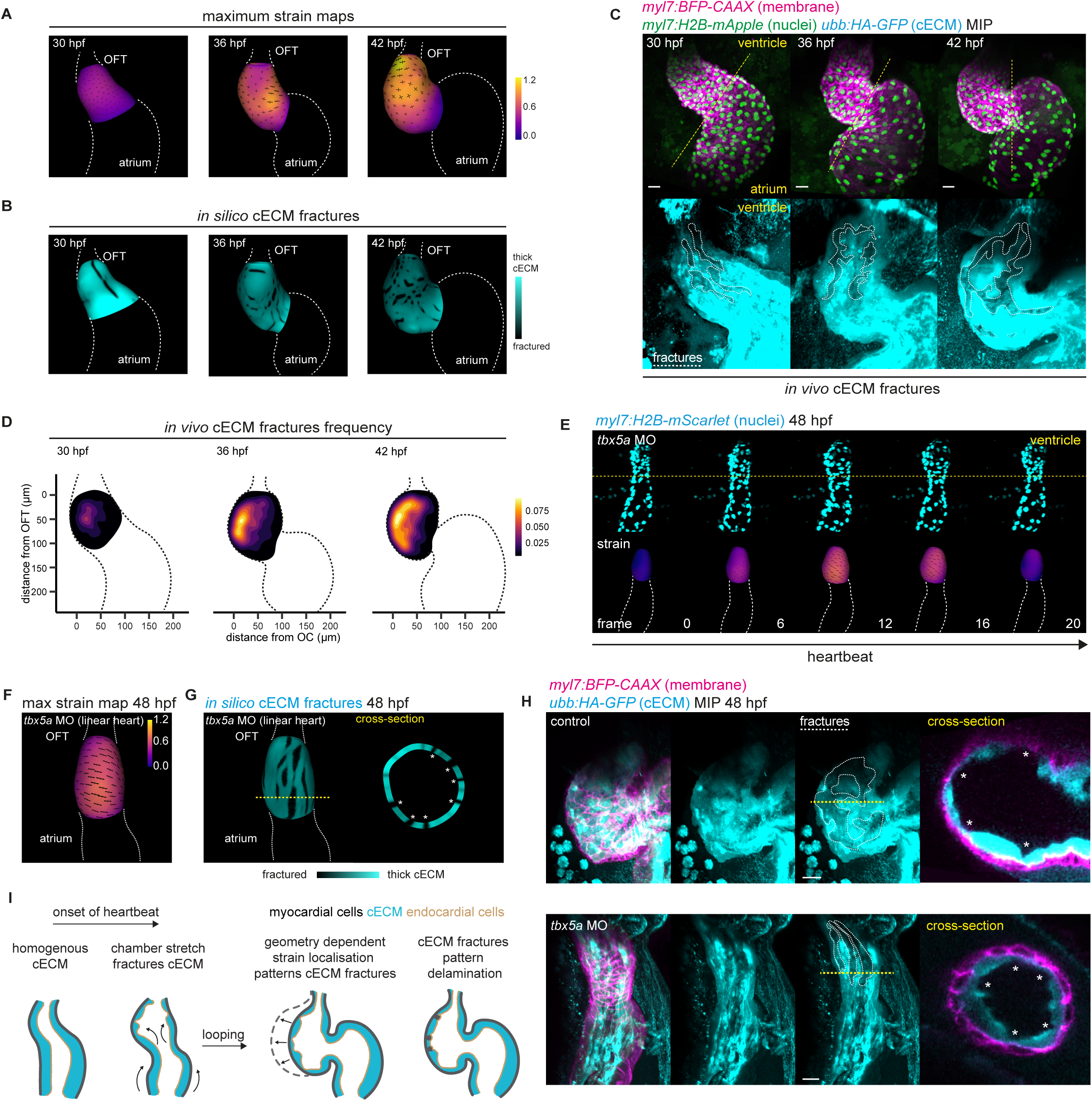
Tissue geometry constrains cECM fractures by pattering mechanical strain. **(A-B)** Maximum myocardial tissue strain maps **(A)** and corresponding *in silico* cECM fractures **(B)** for a 30 hpf, 36 hpf and 42 hpf heart, showing evolution of strain maps and fractures as heart geometry changes. Arrows in **(A)** indicate the directions of maximum and minimum strain, and their size represents the magnitude of the strain; OFT, outflow tract. **(C-D)** Maximum intensity projection (MIP) showing changes in heart geometry and cECM fractures in the ventricle **(C)** and frequency heatmap of cECM fractures distribution in 30 hpf (n = 13), 36 hpf (n = 17), and 42 hpf (n = 13) ventricles, showing changes in patterning of cECM fractures as the heart geometry changes **(D).** *myl7:BFP-CAAX* marks cardiomyocyte membrane, *myl7:H2B-mApple* marks cardiomyocyte nuclei, *ubb:HA-GFP* labels cECM. **(E-F)** Myocardial tissue strain maps calculated across a heart cycle **(E)** and ventral view of maximum strain map of a 48 hpf linear *tbx5a* morphant (MO) ventricle **(F).** *myl7:H2B-mScarlet* marks cardiomyocyte nuclei. Arrows indicate the directions of maximum and minimum strain, and their size represents the magnitude of the strain; OFT, outflow tract. **(G)** *in silico* cECM fractures pattern of a 48 hpf *tbx5a* morphant ventricle showing linear fractures in ventral view and evenly spaced fractures in cross-section view; OFT, outflow tract. **(H)** Patterning of cECM fractures in 48 hpf control (n = 21/21 embryos) and *tbx5a* morphant ventricles (n = 17/23 embryos) *in vivo* showing linear cECM fractures in ventral view and evenly spaced fractures in cross-section view. **(I)** Graphical summary: at the onset of heart contraction, mechanical strain arising from the myocardial tissue stretching initiate cECM fracturing. As the heart loops, changes in myocardial geometry leads to the highest strain in the outer curvature, which in turn spatially confines cECM fractures, thereby patterning delamination. All scale bars are 20µm.

Together, these findings suggest that myocardial tissue geometry influences the patterning of cECM fractures. To further strengthen this notion, we asked whether altering tissue geometry is sufficient to modify the pattern of cECM fractures. We manipulated heart geometry *in vivo* using a well characterized *tbx5a* morpholino, which disrupts heart looping thus generating linear hearts (*11*) (**Fig. 4E**). Indeed, changing tissue geometry is sufficient to change the deformation pattern. In these 48 hpf linear hearts, the direction of maximum strain is more homogenous compared to wild type 48 hpf hearts (**Fig. 3G**) and lies perpendicular to the long axis of the heart (**Fig. 4E-F, Supplementary Video 13**). When the deformation dynamics of linearized hearts were applied to the cECM in simulations, fractures align perpendicular to the maximum principal strain and run linearly across the heart (**Fig. 4G, Supplementary Video 14**). Remarkably, consistent with our modelling prediction, *in vivo* analysis of *tbx5a* morphant hearts revealed predominantly linear fractures running along the long axis of the heart (**Fig. 4H**). Further, the fracture pattern appears less biased and more evenly distributed across the chamber in a cross-section, in line with our modelling predictions. (**Fig. 4H**). We further confirmed these findings by generating linear hearts by inhibiting BMP signalling with K02288, as performed previously (*14*) and observed similar linear cECM fracture patterns (**Fig. S8F**). Overall, we conclude that myocardial tissue geometry instructs its deformation dynamics, which in turn, governs the patterning of cECM fractures.

Overall, we demonstrate that the interplay between mechanical cECM fracturing and tissue geometry guides myocardial tissue patterning (**Fig. 4I**). Our quantitative analysis of Nrg2a-Erbb2 signalling reveal no enrichment in the outer curvature, thus arguing that previously proposed mechanisms of biochemical pre-patterning is not sufficient to explain biased delamination in the outer curvature (*13, 16-18, 20-22, 25, 27-30*). Instead, we find that mechanical patterning of cECM fractures patterns delamination in the outer curvature.

While previous studies have shown that enzymatic activity regulates cECM dynamics during heart morphogenesis (*34, 35, 40, 41*); in this study, integrating 3D morphometrics, mechanical stress mapping, cross-species comparison, predictive modelling and controlled perturbations, we demonstrate that mechanical forces generated by heart contraction is also sufficient to fracture the cECM. Specifically, higher mechanical strain in the outer curvature generates and patterns mechanical fractures that facilitate delamination at the onset of trabeculation. These cECM fractures could possibly facilitate delamination by promoting inter-tissue interaction through endocardial protrusions, as postulated before (*20, 21*). In mouse, fine-tuning of cardiac jelly (cECM) dynamics by Notch and neuregulin1 signalling drives endocardial sprouting to form dome-like matrix-rich structures, which is important for later stages of trabeculae growth and architecture (*18, 26*). Whether cECM remodelling similarly promotes trabeculae growth at later stages in zebrafish remains to be studied.

Living materials are constantly challenged with dynamic mechanical deformations. Accordingly, they adapt to resist any damage or fracture, as fractures compromise biological barriers or lead to disease conditions (*70*). Here, we demonstrate that mechanical fracturing of the cECM is actually utilized as a morphogenetic tool to shape the developing myocardial tissue. Unlike the chaotic dynamics of fracturing in inert materials (*55, 71, 72*), these cECM fractures follow a spatial pattern defined by tissue geometry, thereby robustly patterning morphogenesis. Our findings strengthen the emerging role of fractures in sculpting tissues (*73-79*). Further, we offer key insight into how living materials adapt or yield to extreme dynamical forces at faster timescales (milliseconds), which is a prominent area of focus in the field of material science (*80*) paving the way for innovative biomimetic designs.

It is increasingly evident that the extracellular matrix is not a passive boundary but undergoes dynamic mechanical remodelling to drive morphogenetic changes during development (*36, 37, 81-83*) or cancer invasion (*84-87*). While most research has focused on basement membrane remodelling (*38, 81, 83, 88, 89*), we now provide compelling evidence that hyaluronan matrices also undergo mechanical remodelling. This supports the growing notion that the physical properties of hyaluronan matrices can be modulated to shape diverse tissues during development (*90-93*). Given that hyaluronan is abundant during embryonic development (*94-96*) and many developing tissues generate dynamic mechanical forces as part of their normal physiology (e.g., gut and lungs), it is plausible that mechanical remodelling of hyaluronan matrices is a more widespread mechanism than previously appreciated.

Dynamic changes in tissue geometry are a hallmark of morphogenesis. While much emphasis has been put on how complex geometrical orders (e.g. folds and curvature) emerge, how tissue geometry feeds back to morphogenesis remains less understood (*4, 5, 63*). Here, we show that tissue geometry is not only a consequence of morphogenesis but can also constrain the outcome of morphogenesis by patterning mechanical stress. Our findings emphasize the self-regulating nature of embryonic morphogenesis, where tissue boundary conditions defined by its geometry or ECM, act as a template to steer the course of its own patterning.

Collectively, our work provides key insight into geometric and ECM control of robust tissue pattering in a developing, functioning organ. Tissue geometry and mechanical ECM remodelling are also known to influence cellular processes in disease contexts (*97, 98*). Further, imposing geometrical or ECM constraints lead to reproducible physiological patterning in organoids and gastruloids (*99-101*). We thus believe our findings will have fundamental implications for organ biology, regenerative biology and synthetic morphogenesis.

## Supporting information

Chan et al-supplementary modelling

## Author contributions

C.C.J.J. and R.P. conceived the study. C.C.J.J. designed and performed most of the experiments, conceived the methods for early heart imaging, and analysed, visualised and curated the data. D.S-O. and A.T-S. conceived, designed and performed the strain inference methods and computational modelling, with conceptual input from C.C.J.J. and R.P. M-C.R. generated the *myl7:H2B-mApple^fci604^* line, performed all HCRs with C.C.J.J., and helped with protocol optimization and troubleshooting. J.S-P. and E.N. conceived and developed morphoHeart, and helped apply it to the lines used in the project. T.A. generated the heatmap, 2D cECM and ERK quantification methods. P.P. generated in situ hybridisation probe constructs, and J.S.P performed all in situ hybridisations. C.C.J.J., R.P., D.S-O., and A.T-S. interpreted the data. C.C.J.J. and R.P. wrote the manuscript with input from all other authors. R.P. supervised the overall project, and A.T-S. supervised the modelling part of the project.

## Acknowledgements

We thank all members of the Priya Lab for helpful discussions throughout the project. We are grateful to Alpha Yap, James Sharpe, Yanlan Mao, Caroline Hill, Alberto Elosegui-Artola, Stefan Harmansa, Vikas Trivedi, Talya Dayton and Jim Swoger for critical reading of the manuscript. Special thanks to M-C. Ramel for her expertise in zebrafish handling and techniques, as well as for her invaluable help in optimizing and testing various antibodies and constructs. We thank Giulia Boezio for her guidance on chick husbandry, Srinivas Allanki for his input on mechanobiology techniques, Louise Carroll for optimizing HCR protocols, Alessandra Gentile and Felix Gunawan for initial discussions at the outset of this project. We thank T. Pockert, K. Stoneman, G. Delaqua, M. Millington and the rest of the Crick aquatics team for their support with zebrafish husbandry, and K. Anderson and D. Gunton from the Crick Advanced Light Microscopy facility for support and guidance with imaging techniques. We thank Marco Musy for his support with vedo. A.T-S. and D.S-O are supported by the European Molecular Biology Laboratory (EMBL). J.S-P. was supported by BBSRC Standard Grant BB/W004305/1. E.N. was supported by a British Heart Foundation Fellowship award FS/16/37/32347. Work in R.P.’s laboratory is supported by the Francis Crick Institute, which receives its core funding from the Cancer Research UK (FC011160), the UK Medical Research Council (FC011160), the Wellcome Trust (FC011160), and the British Heart Foundation (SP/F/20/ 150014).

## Supplementary Figure Legends

**Supplementary Fig. 1: Nrg2a-Erbb2 signalling is not enriched in the outer curvature of the ventricle.**

**(A)** Image processing pipeline for quantifying single cell behaviour in 3D. Nuclei were segmented using Imaris, and coordinates were extracted, normalized and pooled to generate an average ventricle shape. Cells of interest were identified and pooled to generate a frequency heatmap.

**(B)** Image processing pipeline for quantifying localized gene expression with hybridization chain reaction (HCR). Raw HCR signal attenuated with FIJI; nuclei were segmented using Imaris, and classified according to their location; outer curvature (OC); ventral face (VF); inner curvature (IC)

**(C)** *erbb2* expression in 48 hpf ventricles analysed using HCR showing no enrichment in the outer curvature; midsagittal sections (top), maximum intensity projection (MIP, bottom), and associated quantification; OC, n = 80 cells; VF, n = 134 cells from 11 embryos. *myl7:BFP-CAAX* marks cardiomyocyte membrane, *kdrl:NLS-mCherry* marks endocardium nuclei.

**(D)** *nrg2a* expression in 48 hpf ventricles analysed using HCR showing no enrichment in the outer curvature; midsagittal sections (top), maximum intensity projection (MIP, bottom) and associated quantification, OC, n = 123 cells; VF, n = 194 cells from 15 embryos.

**(E)** Image processing pipeline for quantifying ERK activity based on the nuclear to cytoplasmic ratio of KTR-Clover fluorescence intensity. KTR-Clover is transported out of the nucleus when ERK is active. Nuclear and peripheral cytoplasmic masks were segmented using FIJI; ERK activity is defined as fluorescence intensity ratio of less than 1.15.

**(F-G)** Representative MIP showing KTR-Clover signal (ERK activity) in the 48 hpf ventricle **(F),** frequency heatmap of ERK active cells (outflow tract, OFT; outer curvature, OC); cyan box, OC. n = 21 embryos **(G).** KTR-Clover is transported out of the nucleus when ERK is active. White dotted region indicates area where signal intensity is too weak for an accurate ratiometric analysis.

All data are mean ± s.d. P values result from unpaired two-tailed Students’ t-tests. Scale bars, 20µm.

**Supplementary Fig. 2: Heterogeneities in cECM thickness, and not basement membrane, patterns delamination in the outer curvature.**

**(A-E)** Fibronectin and Laminin localization in 48 hpf ventricle, n = 26/26 and 22/22 embryos respectively **(A, B)** and 55 hpf ventricle, n = 10/10 and 13/13 embryos respectively **(C, D),** normalized mean fluorescence intensity of Laminin over CL and DL cells, n = 27 cells from 13 embryos **(E).**

**(F)** Midsagittal sections of a 48 hpf ventricle showing cECM marked by *ubb:HA-GFP* sandwiched between myocardium (*myl7:BFP-CAAX*) and endocardium (*kdrl:HRAS-mCherry*).

**(G)** Midsagittal section of a 48 hpf ventricle and corresponding 2D cECM thickness maps from the marked area (white dotted line) showing heterogeneity in cECM thickness before delamination.

**(H)** Midsagittal sections of a 48 hpf ventricle with corresponding 2D cECM thickness map from the marked area (dotted line); cECM thickness segmented from negative space between the endocardium (*kdrl:lifeact-GFP)* and myocardium showing heterogeneity in cECM thickness before delamination.

**(I)** Midsagittal sections of a 55 hpf ventricle with corresponding 2D cECM thickness map from the marked area (dotted line); cECM thickness segmented from negative space between the endocardium and myocardium showing heterogeneity in cECM thickness at the onset of delamination

**(J)** Schematic showing segmentation pipeline of morphoHeart. For more details, please see Methods.

**(K-L)** Quantifications showing that cECM fractures are independent of cell size **(K)** and cell shape (circularity index) **(L)**, thick cECM n = 134 cells, cECM fractures n = 87 cells from 35 embryos

**(M)** MIP from longitudinal imaging of the same ventricle at 48 hpf and 55 hpf with one cell mosaically labelled with mCardinal, interacting with the surrounding cECM fractures by extending protrusions. mCardinal cells with protrusions in fractures observed in n = 8/11 embryos at 55 hpf.

**(N-P)** Midsagittal section of a 72 hpf ventricles overexpressing mApple and Vcanb **(N)** quantification of cECM thickness, mApple (n = 75 cells), Vcanb (n = 44 cells) **(O)** and delamination frequency, mApple (n =12 embryos) and Vcanb (n = 13 embryos) **(P)**.

Data are mean ± s.d. P values result from unpaired two-tailed Students’ t-tests. Scale bars, 20µm.

**Supplementary Fig. 3 Nrg2a-Erbb2 activity and Notch activity is not required to fracture the cECM.**

**(A-B)** 48 hpf wild type and *nrg2a* mutant ventricle, representative of 16/16 and 18/18 embryos respectively, representative 3D cECM thickness heatmap showing no significant changes in cECM distribution **(A)** and average 2D cECM thickness heatmaps generated with morphoHeart **(B)** wild type sibling, n = 7 embryos, *nrg2a* mutant, n = 6 embryos.

**(C)** 55 hpf ventricles from DMSO (control) and PD168393 treated embryos showing no significant changes in cECM distribution, representative of 17/17 embryos and 15/15 embryos respectively. White asterisks, delamination.

**(D-E)** 48 hpf ventricles from DMSO (control) and LY411575 treated embryos showing no significant changes in cECM distribution, representative of 10/10 embryos each, representative 3D cECM thickness heatmap **(D)** and average 2D cECM thickness heatmaps generated with morphoheart **(E)** DMSO, n = 5 embryos, LY411575 treated, n = 5 embryos

**(F)** 55 hpf wild type and *nrg2a* mutant ventricles, representative of 16/16 embryos and 18/18 embryos respectively. White asterisks, delamination.

Scale bars, 20µm.

**Supplementary Fig. 4 cECM fractures emerge independent of enzymatic activity**

**(A)** 48 hpf ventricles from DMSO (control) and GM6001 treated embryos, representative of 10/10 embryos each, and corresponding 2D cECM thickness map from the marked area (white dotted line).

**(B)** *in situ* hybridisation expression patterns of hyaluronidases at 26 hpf and 50 hpf. White arrows indicate enrichment of signal in the ventricle. Representative numbers as indicated.

**(C-E)** *tmem2* expression in 48 hpf ventricles **(C)** quantified in the outer curvature (OC) vs. ventral face (VF) **(D)** or thick vs. cECM fractures (frac) **(E)** in myocardial and endocardial layers using hybridization chain reaction (HCR) showing no biased expression pattern. Normalized sum *tmem2* intensity in outer curvature (OC); n = 49 endocardial cells (endo), n = 75 myocardial cells (myo) versus ventral face (VF); n = 70 endo, n = 84 myo from 11 embryos

**(D)** normalized sum *tmem2* intensity in thick cECM; n = 70 endo, n = 64 myo, versus cECM fractures (frac); n = 72 endo, n = 67 myo from 11 embryos **(E).**

**(F-H)** *hyal2b* expression in 48 hpf ventricles **(F)** quantified in the outer curvature (OC) vs. ventral face (VF) **(G)** or thick vs. cECM fractures (frac) **(H)** in myocardial and endocardial layers using hybridization chain reaction (HCR) showing no biased expression pattern. Normalized sum *hyal2b* intensity in outer curvature (OC); n = 47 endo, n = 48 myo versus Ventral face (VF), n = 55 endo, n = 59 myo from 11 embryos **(G)** Normalized sum *hyal2b* intensity in thick cECM; n = 42 endocardial cells (endo), n = 40 myocardial cells (myo) versus cECM fractures (frac); n = 42 endo, n = 40 myo from 11 embryos **(H).**

**(I)** *has2* expression in 48 hpf ventricles analysed using hybridization chain reaction (HCR) showing enrichment in the valve region. n = 19/19 embryos.

Data are mean ± s.d. P values result from unpaired two-tailed Students’ t-tests. Scale bars, 20µm.

**Supplementary Fig. 5 cECM fracturing is driven by myocardial tissue contractility independent of shear stress.**

**(A-B)** Schematic showing the surgical removal of the anterior portion of the embryo to make early-stage hearts optically accessible (body explant method) **(A)**, whole mount images of 24 hpf body explants (BE) and sibling control embryos, linked with Supplementary video 3 **(B)**.

**(C)** Maximum intensity projections of cECM in 30 hpf ventricles, representative of 15/17 embryos and corresponding zoomed view from the marked area (dotted box) showing linear tissue-scale fractures.

**(D)** 48 hpf ventricles from DMSO (control) and Nifedipine treated embryos, representative of 13/13 embryos each and corresponding 2D cECM thickness map from the marked area (white dotted line).

**(E)** Distribution of cECM in 48 hpf wild type (wt) and *half hearted (haf)* mutant ventricles, representative of 13/13 embryos each and corresponding 2D thickness map from the marked areas (dotted lines).

**(F-G)** 48 hpf ventricles from control and *gata1* morphant (MO) embryos, representative of 26/26 and 23/23 embryos respectively, corresponding 3D cECM thickness heatmap **(F)** and average 2D cECM heatmaps generated with morphoHeart **(G)** control, n = 5 embryos, *gata1* morphant, n = 6 embryos

Scale bars, 20 µm.

**Supplementary Fig. 6 cECM fractures also exist in chick embryonic hearts and are driven by myocardial tissue contraction.**

**(A)** cECM distribution in early chick hearts analysed using negative space between myocardium and endocardium labelled with cytoplasmic GFP at HH10 showing homogenous distribution before the heart starts beating, representative of n = 7/8 embryos, and at HH12 showing fractures at the onset of heart contraction, representative of n = 8/8 embryos.

**(B)** cECM distribution in HH12 DMSO (control), representative of n = 10/10 embryos, and Nifedipine treated chick embryos, representative of n = 11/11 embryos showing that abrogation of heart contractility affects fractures.

Scale bars, 20 µm.

**Supplementary Fig. 7 cECM fractures are mechanically driven by myocardial tissue contraction.**

**(A)** Schematic showing how nuclei are tracked in Imaris to generate nuclear coordinates across a heartbeat.

**(B)** Schematic showing how nuclei track data is used to reconstruct myocardial deformation. A three-dimensional reference surface is reconstructed using the screened Poisson surface reconstruction algorithm from the reference position of the nuclei. The deformation map is then obtained by interpolating and smoothing the displacement of the nuclei. Strain maps are obtained from the right-Cauchy deformation tensor obtained from the deformation map.

**(C)** Distribution of cECM in 48 hpf ventricle after being stopped for 2.5 hrs showing that it does not dissipate, n = 26 embryos.

**(D)** cECM morphology in decellularized 48 hpf hearts showing that it retains its shape and behaves like an elastic solid, n = 8 embryos.

**(E)** MIP and FSI (flattened sum intensity) of the same ventricle analysed at 36 hpf and 42 hpf showing that initial fractures deepen and become prominent over time, n = 8 embryos.

**(F-G)** cECM distribution in a 48 hpf ventricle treated with DMSO (control, n = 10 embryos), IBMX (n = 9 embryos), Isoprenaline (n = 9 embryos), Nifedipine (n = 10 embryos) and Lidocaine (n = 10 embryos), corresponding 3D cECM thickness heatmap **(E)** and average 2D cECM thickness heatmap **(F)** derived using morphoHeart, n = 6 embryos each

**(H)** Quantification of heart rate in control vs *amhc* morphant (MO) hearts showing *amhc* morphant MO ventricle does not beat slower, n = 15 embryos each

**(I)** Nuclei positions and corresponding area strain across a heart cycle in an *amhc* MO 48 hpf ventricle, *myl7:H2B-mScarlet* marks the nuclei of cardiomyocytes

**(J-K)** Analysis of delamination in 55 hpf ventricles treated with DMSO (control), IBMX, Isoprenaline, Nifedipine and Lidocaine **(J)** and quantification of number of delaminating cells in drug treated hearts **(K)** control n = 17, IBMX n = 18, Isoprenaline n = 18, Lidocaine n = 8, Nifedipine n = 13 embryos.

Data are mean ± s.d. P values result from unpaired two-tailed Students’ t-tests. Scale bars, 20µm.

**Supplementary Fig. 8 Myocardial tissue geometry patterns cECM fractures.**

**(A)** Mean distribution of *in silico* cECM fractures in 48 hpf ventricles, n = 3.

**(B-D)** Nuclei positions and corresponding area strain across a heart cycle in a 30 hpf, 36 hpf and 42 hpf ventricle. *myl7:H2B-mScarlet* marks the nuclei of cardiomyocytes.

**(E)** Mean distribution of *in silico* cECM fractures in 30 hpf, 36 hpf and 42 hpf ventricles; OFT, outflow tract, n = 3.

**(F)** cECM distribution in 48 hpf DMSO (control, representative of n = 13/13 embryos) and K02288 treated ventricles (BMP inhibition) (representative of n = 9/12 embryos) showing linear cECM fractures in ventral view and evenly spaced fractures in cross-section view;

Scale bars, 20µm.

## Supplementary Video Legends

**Video 1. cECM fractures run extensively across the myocardium in a 48 hpf ventricle**

Video showing the rotating maximum intensity projection of a 48 hpf heart created with Imaris.

**Video 2. Cells delaminate into cECM fractures at 55 hpf**

Video showing the rotating maximum intensity projection of a 55 hpf heart created with Imaris. Delaminating cells pseudo-coloured using the manual surfaces function.

**Video 3. 24 hpf body explants have beating hearts and undergo normal heart development until 48 hpf**

Wholemount timelapse movies of beating hearts and blood flow in 24 hpf body explants and 48 hpf, 24 hours after surgery, acquired at 12.3 fps.

**Video 4. 24 hpf beating heart showing the optical resolution with the body explant method**

Video showing a maximum intensity projection of a beating 24 hpf heart temporally registered with Beatsync, acquired at 100 fps showing cellular resolution.

**Video 5. cECM is homogenous at 24 hpf and fractured at 30 hpf**

Midsagittal timelapse movies of a 24 hpf and 30 hpf heart, acquired at 100 fps.

**Video 6. Nuclei tracks of a beating 48 hpf heart**

Video showing tracked nuclei in a maximum intensity projection of a beating 48 hpf heart, temporally registered with Beatsync. Nuclei were projected and tracked with Imaris, acquired at 100 fps.

**Video 7. Strain maps of a beating 48 hpf heart showing higher strain in the outer curvature**

**Video 8. Simulated cECM fracture dynamics of a beating 48 hpf heart showing fractures emerging in the outer curvature**

**Video 9. Strain maps of a beating 48 hpf *amhc* morphant heart showing lower strain**

**Video 10. Simulated cECM fracture dynamics of a beating 48 hpf *amhc* morphant heart showing less fractures emerging**

**Video 11. Strain maps of beating 30, 36 and 42 hpf hearts showing the shift in strain patterns as ventricle geometry evolves**

**Video 12. Simulated cECM fracture dynamics of beating 30, 36 and 42 hpf hearts showing the shift in cECM fracture patterns as ventricle geometry evolves**

**Video 13. Strain maps of a beating 48 hpf *tbx5a* morphant heart showing homogenous strain aligned perpendicular to the long axis of the heart**

**Video 14. Simulated cECM fracture dynamics of a beating 48 hpf *tbx5a* morphant heart showing linear fractures running along the long axis of the heart**

## Material and Methods

### Zebrafish handling

*Danio rerio* were maintained at 28.0°C in fresh water (pH 7.5 and conductivity 500 µS) on a 15 hour on/9 hour off light cycle, in accordance with institutional (The Francis Crick Institute) and national (United Kingdom) ethical and animal welfare regulations. All procedures were performed under project license PP8356093, in accordance with UK Home Office Regulations, as per the Animals (Scientific Procedures) Act 1986. All zebrafish embryos used for live-imaging experiments were treated with 0.003% phenylthiourea (PTU, Sigma-Aldrich P7629) from 24 hpf to prevent the formation of pigmentation.

### Zebrafish transgenic lines and mutants

The transgenic lines and mutants used in this study and corresponding abbreviations are listed in **Table S1.**

### Plasmid Construction

All plasmids generated in this study for transgenesis used a pTol2 vector backbone that contains Tol2 sites necessary for Tol2-mediated transgenesis (*102, 103*). For cloning, appropriate restriction enzymes were used to linearize the pTol2 vector. DNA fragments to be inserted were amplified by PCR using Phusion DNA polymerase (NEB M0530) with appropriate primers (see **Table S2**), either from cDNA or from synthesised constructs (Twist Bioscience). In-Fusion® technology (Takara Bio #638947) was used to assemble the desired plasmids according to manufacturer’s protocol. All plasmids were sequenced and verified using Sanger sequencing and/or full plasmid sequencing.

### Mosaic Analysis

Tol2 transgenesis was performed as previously described (*102, 103*). Briefly, an injection mix containing Tol2 mRNA (15 pg per embryo) and plasmid DNA (15 pg per embryo) was injected into embryos at the one-cell stage. Embryos were screened at desired stages for mosaic fluorescence prior to confocal imaging.

### Morpholino injections

Morpholinos were injected at the one-cell stage. All morpholinos used in this study have been well characterized and validated in previous studies.

*tnnt2a* morpholino; 5’-CATGTTTGCTCTGATCTGACACGCA-3’ (0.3 ng) (*16, 17, 20, 27, 104*)

*amhc* morpholino 5’-ACTCTGCCATTAAAGCATCACCCAT-3’ (0.2 ng) (*17, 57, 62, 105*)

*gata1* morpholino 5’-CTGCAAGTGTAGTATTGAAGATGTC-3’ (8 ng) (*39, 59, 106*)

*tbx5a* morpholino 5’-GAAAGGTGTCTTCACTGTCCGCCAT-3’ (1.2 ng) (*107, 108*)

*tmem2* morpholino 5’-ACAAACCAAAGCCATCTCACCTTGA-3’ (0.5ng) (*53, 109*)

*has2* morpholino 5’-AGCAGCTCTTTGGAGATGTCCCGTT-3’ (3ng) (*41, 54*)

All morpholino injection mixes were heated to 65°C for 10 minutes in a heat block and cooled to room temperature before injections, as done previously (*110, 111*)

### Zebrafish transgenesis

Tol2 transgenesis was used to generate the *Tg(-0.8myl7:H2B-mApple)fci604* line. Wild type embryos were injected at the one-cell stage with an injection mix of transposase (*Tol2*) mRNA (15 pg per embryo) and plasmid DNA (15 pg per embryo). F0 embryos positive for mApple fluorescence in the myocardium were raised to adulthood before being screened for germline transmission. Adult founders were outcrossed to wild type animals, and their progeny were screened for morphologically healthy hearts with homogenous nuclear mApple fluorescence in the myocardium. At least 3 founders were identified, and the founder with progeny with the healthiest hearts were outcrossed to raise the F_1_ generation.

### Stopped heart imaging

Embryos were screened for positive fluorescence using a Leica SMZ18 microscope. Selected embryos were mounted in 35 mm diameter glass-bottomed dishes (MatTek P35G-1.5-14-C), in 1.2% (w/v) low-melting agarose / E3 medium containing 2 µg/µl tricaine (Merck A5040) to stop the hearts. Once the agarose was set, the dish was filled with 4 µg/µl tricaine. All stopped-heart images were acquired using a Zeiss LSM980 Axio Examiner confocal microscope with a Zeiss W Plan-Apochromat 40x / 1.0 DIC M27 water immersion dipping lens.

Embryos were mounted at different angles depending on the stage. For 24 hpf embryos, embryos were mounted at a 90° angle (tail facing downwards away from the objective), such that the dorsal (future ventral) face of the heart is perpendicular to the light path from the objective. As the embryo develops and the heart descends, embryos were mounted at increasing angles to ensure the ventral face of the heart faces the objectives – 100° for 30 hpf, 110° for 36 – 48hpf and 120° for 55 hpf.

### Longitudinal timelapse imaging

For repeated imaging of the same embryo, embryos were gently released from agarose immediately post-imaging using forceps, and washed once in E3 media, before being isolated in 6 well plates containing 6ml of E3 media. These fish were then returned to a 28.0°C incubator to allow normal development to resume.

### Imaging of zebrafish hearts from 24 hpf to 42 hpf

Due to depth of cardiac tissue at early stages, only the superficially located venous pole/future atrium of the heart tube can be imaged with sufficient resolution (*41*). To image arterial pole/ventricle development, we surgically removed the anterior region (anterior to the otic placode) of embryos with N.5SF Dumont forceps 1 hour before the desired stage. Undamaged posterior regions (identified by an intact yolk sac, undamaged body and beating heart, **Supplementary Video 3**), which we refer to as body explant (BE) fish, were transferred to supplemented L15 medium for an hour (15% fetal bovine serum, 0.8 mM CaCl2, 50 μg/ml penicillin, 0.05 mg/ml streptomycin, 0.05 mg/ml gentamicin), as previously described for ex-vivo heart cultures [10]. Note that L15 media should be made fresh, if possible, but can be stored at -20°C for up to two weeks if necessary. After an hour of recovery, healthy BE fish which show closed systemic circulation, and an inflated beating heart were processed as previously described for either stopped heart or longitudinal timelapse imaging as desired. To confirm that removal of the anterior region of the embryo does not affect heart development, some BE fish were allowed to develop overnight from 24 hpf in E3 media, until 48 hpf. The hearts of these BE fish undergo looping and form two distinct chambers (**Supplementary Video 3**), confirming that the surgery does not abrogate heart development.

### Timelapse imaging of beating hearts

For timelapse imaging of beating hearts, embryos were embedded in 1.2% (w/v) low-melting agarose, with 0.06 µg/µl tricaine to immobilise the embryo with limited effect on heartbeat, in 35 mm diameter glass-bottomed dishes. Embryos were imaged using a Nikon W1 spinning disk confocal microscope with a Nikon Apo LWD 40x WI λS DIC N2 water immersion dipping objective, in a heated chamber kept at 28.0°C. All timelapse imaging was performed with an exposure time of 10 ms. For 3D timelapse imaging of beating hearts, 100 frames were acquired sequentially for each Z-slice over a 150 µm Z-depth, with 1.5 µm slice spacing. Post-acquisition temporal registration, Beatsync, was then used to align imaging data from each slice, to assemble a 4D movie (*112*)

### Image processing

For representation of data in 2D, mid-sagittal images were processed in FIJI by using the smooth function, applying a median filter of 1.2-1.5 pixel radius, as appropriate. When necessary, background subtraction was performed as well, with a rolling-ball radius of 50-100 pixels radius, as appropriate. For representation of data in 3D, signal from the ubiquitously expressed *ubb:HA-GFP* channel was manually pre-processed before projections. Non-specific signal (i.e. signal from hatching glands and pericardial membrane) was manually deleted in FIJI, if necessary, before the signal was projected. This manual deletion was only performed for hearts where non-specific signal interfered with the projection of signal from cECM. For an example of non-specific signal, see **Figure 4H**. Both FIJI and Imaris was used to produce 3D Maximum intensity projections.

### Heatmap analysis of delamination and cECM fracture distribution

Nuclear coordinates from individual ventricles were superimposed using a modified Procrustes superimposition, scaling each axis independently in length (**Supplementary Fig. 1A**). For an average spatial analysis of cellular distributions, nuclei from individual ventricles were first aligned to a standardised orthogonal space, determined using the coordinates of paired ventricle landmarks (Valve – Outer Curvature, Apex – Outflow track, Dorsal – Ventral), as determined in Imaris. The 3D coordinates of the landmarks and nuclei were exported, and an R script was used to infer a new vector of points running between (and extrapolating beyond) each pair of landmarks, separated by 1 µm. The position of each nucleus along each axis was then calculated by selecting a point on the axis that lies closest to the nucleus (shortest Euclidean distance).

To aid superimposition, the length of each axis was adjusted to the mean length of the whole population. For this, the maximum distance between nuclei along each axis was determined and normalized between 0 and 1 across the whole population, before being multiplied by the mean of the whole population. This allows each ventricle to scale freely on each axis.

All coordinates from the pooled nuclei were then plotted in a normalised coordinate system, using the stat_density2d function in the ggplot2 package to generate heatmaps for each cell state. A first layer was generated utilizing all nuclear coordinates, representing mean ventricle shape. A second layer was then added depending on the feature of interest, in the case of this study either delaminating cells, ERK positive cells, or cells that lie in cECM fractures. This generates a heatmap showing the spatial distribution of these cells in a ventral view.

Delaminating cells were identified as having a constricted apical domain (*17*). Cells that lie in cECM fractures were determined in Imaris, where fractures were defined as regions of cECM where the thickness was less than 2µm. All delaminating cells and cells that lie in cECM fractures were manually classified and analysed across three orthogonal axes.

### 2D#cECM thickness graphs

2D cECM thickness graphs were generated to quantitatively represent the heterogeneity of the cECM in representative regions of the heart (for example, see **Fig 1D, 1J**), and to quantify the thickness above delaminating versus compact layer cells. Briefly, the region of interest was first straightened in FIJI, before a Gaussian Blur of radius 2 was applied. The signal was then converted to a binary mask using the Threshold function, and any signal that was not part of the cECM was subsequently manually deleted. The mask is then saved as a text image, before being imported into a custom R script. Briefly, the script counts the number of positive binary events (indicating that signal is present) in every column, before multiplying that value by the pixel size to determine the thickness of the cECM.

For 2D cECM graphs determined from the negative space between the myocardium and endocardium, after a region of interest was straightened in FIJI, signal from both tissue layers were combined in the same channel, and a Gaussian Blur of radius 2 was applied. The signal was then converted to a binary mask using the Threshold function, and the binary values were inverted using the Invert function. Signal that is not part of the cECM is deleted, and the mask is then imported into the same R script as described above.

### Heatmap analysis of ERK activity

*myl7:KTR-Clover* embryos were used to analyse the distribution of active ERK cells in the heart. When ERK is active, KTR is phosphorylated and transported out of the nucleus (*33*). Using a custom script, a mask of nuclei and a mask of the cytoplasm surrounding the nuclei was created in FIJI. Briefly, nuclei were masked using the Threshold function in FIJI, and the mask was then repeatedly dilated to expand it isotropically. The original nuclear mask is then subtracted from the dilated mask, to yield a mask that represents the cytoplasm that surrounds the nucleus (**Supplementary Fig. 1E)**. The ratio of KTR-Clover intensity between the nucleus and the surrounding cytoplasm was calculated to determine if a cell has active ERK. To exclude cells that have an equal amount of KTR signal in the nucleus and cytoplasm, a ratio of 1.15 was set as the threshold in the analysis performed, as this minimises false positives. Due to depth of cardiac tissue, signal near the outflow tract of the heart was not bright enough for ratiometric analysis and was hence excluded. However, this does not affect our interpretations.

### Ventricle unwrapping

Myocardium membrane signal was unwrapped from the native 3D curvature of the ventricle onto a 2D plane for accurate measurements of cell area and cell circularity, as well as visualization purposes (for example, see **Fig. 1I**). Z-stacks of myocardial signal were rotated in FIJI, without interpolation, to align the anteroposterior axis of the ventricle to the XY axis of the image. The image was then resliced from the left and rotated to align along the ZY plane. Once aligned, Z stacks were resliced from the left to provide a transversal sectioning through the ventricle. A maximum intensity projection was then generated for the middle 50% of the stack, excluding the outflow track and apex, as the middle 50% best approximates to a cylinder. A spline was fit through the centre of the curved signal, and spline thickness was increased to 100 pixels, which was sufficient to include both myocardial and cECM signal. This spline was then applied to the full transversal Z-stack, which was unwrapped using the straighten tool in FIJI. To return to an XY view, the straightened stack was resliced from the top, before being used to generate maximum intensity projections for downstream analysis and visualisation. Flattened Sum intensity projections were used for early-stage embryos (for example, see **Supplementary Fig. 7E**) as the intensity of the biosensor is weaker at these stages.

### Chemical treatments

To stop delamination, embryos were treated with 12 µM of Erbb2 inhibitor PD168393 (*16, 17, 20, 21, 27, 29*) (Merck Life Science #513033) from 34hpf until the desired stage for analysis. To completely stop the heart from beating, embryos were treated with 80 µM Nifedipine (*56*) (Sigma-Aldrich #N7634) from 24hpf, until the desired stage for analysis. To inhibit MMPs, embryos were treated with 100 µM broad-MMP inhibitor (GM6001, Abcam #ab120845) from 24hpf, until the desired stage for analysis. To inhibit heart looping via inhibition of BMP signalling, embryos were treated with 40 µM K02288 (*14*) (Sigma-Aldrich #SML1307) from 30hpf, until the desired stage for analysis. To inhibit Notch signalling, embryos were treated with 2.5 µM of LY411575 (*17*) (Selleck Chemicals #S271) from 30hpf, until the desired stage for analysis. To increase heart rate, embryos were treated with either 50 µM Isobutylmethylxanthane (IBMX; Sigma-Aldrich #I5879) or 100 µM Isoprenaline (Sigma-Aldrich #I5627) (*56*). To decrease heart rate, embryos were treated with either 25 µM Nifedipine (Sigma-Aldrich #N7634) or 0.05% Lidocaine (Merck #PHR1034) (*56*). Embryos were treated from 30hpf until the desired stage for analysis. All the above drugs were dissolved in DMSO to make stock solutions. For all drug treatments, embryos were first incubated in 50 ml petri dishes with 20ml of PTU-treated E3 media with 200 µl of 10 mg/ml pronase for 20 minutes at the desired stage. Dechorionated embryos were then incubated in 50 ml petri dishes with the appropriate concentration of drug in 20 ml of PTU-treated E3 media with 1% DMSO to aid drug solubilization. 1% DMSO was used as a vehicle control for all experiments

### cECM dissipation assay

To analyse if heterogeneities in cECM thickness dissipate over time, hearts were processed for stopped-heart imaging as described above. After imaging, embryos were not recovered from the agarose. Instead, embryos were kept mounted in 35 mm diameter glass-bottomed dishes with 4 µg/µl tricaine, and were incubated at 28.0°C for 2.5 hours, before being imaged again.

### Decellularization of the heart

Hearts were dissected from anesthetised 48 hpf *myl7:BFP-CAAX; ubb:HA-GFP* embryos and immediately placed in a 50 ml petri dish containing 20 ml of supplemented L15 medium (15% fetal bovine serum, 0.8 mM CaCl2, 50 μg/ml penicillin, 0.05 mg/ml streptomycin, 0.05 mg/ml gentamicin) as described above. 5 ml of 4 µg/µl Tricaine was added to the dish to stop the hearts, before the hearts were imaged on the Nikon SMZ18 stereomicroscope. To decellularize the heart, hearts were transferred to 35 mm diameter glass-bottomed dishes containing 4ml of PBS with 1% (w/v) SDS and 1% (v/v) Triton X-100, and swirled gently for 5 minutes. The decellularized heart was then immediately imaged on the Nikon SMZ18, with a Plan APO 1x, W.D.60mm lens. Care was taken to not decellularize for more than 5 minutes as the hyaluronan biosensor does not survive the SDS/Triton treatment.

### mRNA *in situ* hybridization

Embryos were fixed overnight in 4% PFA, and mRNA *in situ* hybridization was carried out as previously described (*15*). Primers used to generate the probes used can be found in **Table S3.**

### Whole-mount Immunostaining

48 hpf and 55 hpf embryos were anesthetized with 4 µg/µl tricaine before fixation. Embryos were fixed at room temperature in 4% paraformaldehyde, 4% w/v sucrose and 120 µM CaCl_2_ in PBS (adjusted to pH 7.35) for 3 hours. Fixed embryos were washed three times using PBST (0.1% Tween-20/1X PBS), and the yolk and pericardial membrane was removed manually with forceps. Embryos were then permeabilized by incubating in 3 µg/ml proteinase-K (Sigma-Aldrich #P6556) dissolved in PBST for 25 minutes. Permeabilized embryos were then washed with PBDT buffer (PBS, 1% BSA, 1% DMSO, 0.5% Triton X-100) 3 times for 10 minutes each, before being incubated in a blocking buffer containing 10% BSA in PBST for 1 hr. Embryos were incubated with primary antibodies at 4°C, on a nutator in PBST buffer, overnight. The next day, embryos were washed three times for 10 minutes each with PBDT, before being incubated in secondary antibody in PBST buffer for 3 hrs at room temperature. This step was followed by a brief 15 minute incubation with DAPI (1:1,000) and five 10-minute washes with PBST.

Primary antibodies used are as follows: Laminin (Sigma-Aldrich #L9393, 1:200), Fibronectin (Sigma-Aldrich #F3648, 1:200), GFP (Aves Labs #gfp-1020, 1:500). Secondary antibodies used are as follows: goat anti-chicken Alexa Fluor 488 (Thermo Fisher Scientific #A-11039, 1:500), goat anti-rabbit Alexa Fluor 555 (Thermo Fisher Scientific #A-21428, 1:500). Nanobodies used are as follows: BFP nanobody (2BScientific #N0502-AF647-L; 1:200), GFP nanobody (2BScientific #N0301-AT488-L-SY; 1:200), mCherry nanobody (2BScientific #N1302-AF568-L-SY, 1:200).

### Whole-mount immuno-coupled Hybridisation Chain Reaction (HCR)

We modified a published HCR protocol (*31*) for analysing gene expression in developing zebrafish hearts. Staged transgenic embryos were first fixed (as above) for 2 hours at room temperature then deyolked in PBST. Deyolking included removal of the yolk and the pericardial membrane above the heart using fine forceps as above. Embryos were then dehydrated gradually in graded methanol/PBS (30%, 50%, 75%, 100%), incubating for 5 min at room temperature at each step and storing at –20°C overnight (they can be stored at –20°C for up to 2 weeks).

Before proceeding with HCR, embryos were rehydrated in graded methanol/PBS (100%, 75%, 50%, 30%, 0), incubating for 5 min at room temperature at each step before being permeabilised overnight with 0.01% DMSO/ 1% Triton. Embryos were then briefly fixed and washed with PBST before proceeding with probe hybridisation overnight (probes diluted 1:50-1:125 in hybridisation buffer).

The following day, embryos were washed extensively with wash buffer followed by 5X SSCT before incubation in amplification buffer. Hairpins were heated for 90 secs at 95°C, and allowed to cool for 3 minutes at room temperature. Embryos were incubated overnight in the hairpin solution, diluted (1:50) in amplification buffer. Nanobodies (1:125) or primary antibodies were added at this stage as needed. The next day, embryos were washed extensively with 5x SSCT. DAPI staining was performed as for whole-mount immunostaining (see above). Hybridisation buffer, wash and amplification buffers were supplied by Molecular Instruments. Detailed protocol available upon request.

Probes targeting *has2, tmem2* and *hyal2b* were ordered from Molecular Instruments. Probes against *nrg2a* and *erbb2* were designed with an in-house script, and the specificity for the *nrg2a* probe was verified in a mutant background. Cardiomyocyte membranes were labelled using a nanobody against BFP (for *myl7:BFP-CAAX*; 2BScientific #N0502-AF647-L, 1:125) and endocardial nuclei were labelled using a nanobody against mCherry (for *kdrl:NLS-mCherry*; 2BScientific #N1302-AF568-L-SY, 1:125).

### Analysis of HCR signal

Due to depth of the tissue and loss of signal at deeper Z stacks, the Attenuation Correction plugin in FIJI was used before HCR signal was quantified, using the first slice where signal can be observed in each stack as the reference slice, with an opening radius of 0.1 µm. This plugin applies a linear transformation of grey levels to each slice image, making the average and standard-deviation of the background constant throughout the stack (*113*). Attenuated images were subsequently processed in Imaris. The Surface wizard was used to generate surfaces of all nuclei, using a detail level of 1 µm. The Split Objects function, with an expected object diameter of 4 µm was then used to subdivide the individual nuclei. Segmented nuclei were assigned to the myocardium or endocardium either manually, or semi-automatically if a tissue-specific nuclear stain was available, by applying an appropriate signal intensity filter. myocardium and endocardium nuclei were then further categorized into outer curvature and ventral face following their position along the Left-Right axis of the heart. Inner curvature nuclei were excluded from analysis as it contains many valve nuclei, which are morphologically and genetically distinct from the rest of the chamber (*114*). The sum HCR signal intensity of each nuclei was calculated using the Statistics function. Myocardium and endocardium nuclei were also categorised into nuclei found in regions of thick cECM or cECM fractures. This was determined by analysing the thickness of cECM in contact with the respective cell in all orthogonal slices. cECM fractures were defined as regions where cECM thickness was less than 2 µm.

### morphoHeart

To produce 3D representative and 2D average thickness heatmaps of the cECM, we utilized morphoHeart software (*46*). Briefly, morphoHeart was used to semi-automatically filter contours to create a 3D volumetric reconstruction. Using confocal stacks of *myl7:BFP-CAAX; ubb:HA-GFP* embryos, the segmented myocardial membrane was used to reconstruct the total myocardial volume and inner myocardium lumen. Similarly, the segmented cECM was used to reconstruct the total cECM volume. As the hyaluronan sensor is under a ubiquitous promoter, the thickness of the cECM cannot be segmented from the channel alone. Hence, the clean function in morphoHeart was used to filter the cECM volume using the myocardial mask, yielding the cECM that is found exclusively within the myocardium (**Supplementary Fig 2J**). As described in morphoHeart, the centreline function was then used to unfold the heatmaps from 3D into 2D, and 2D heatmaps were then superimposed to generate average 2D thickness maps. At least 5 embryos across 2 independent repeats were used for all morphoHeart analyses. Only the ventral half of the heart was analysed in all conditions, as the dorsal half cannot be reliably segmented due to imaging depth.

### Cardiac Physiology measurements

24 hpf, 27 hpf, 30 hpf and 33 hpf hearts were imaged on the Nikon W1 spinning disk confocal microscope with a Nikon Apo LWD 40x WI λS DIC N2 water immersion dipping objective, as previously described for the timelapse imaging of beating hearts. Representative 2D slice movies of beating hearts were analysed in FIJI, and dynamic changes in ventricle width was measured. To calculate heart rate in beats per minute (bpm), the total amount of time taken (in seconds) for 3 beats was determined by multiplying the number of frames required for 3 beats to the known frames-per-second. Dividing this value by 3 yields the number of seconds taken for a beat. As there are 60 seconds in a minute, dividing 60 by this value yields the heart rate in bpm (**Fig. 3C**). Early stage tubular hearts beat in a peristaltic-like motion (*115*) and hence percentage stretch was determined from changes in chamber width, dividing the end-diastolic width by the end-systolic width (**Fig. 3D**).

### Nuclei tracking for strain inference

Nuclei positions across heart beats were semi-automatically tracked using the track function in Imaris. Briefly, 3D timelapse images of beating hearts were acquired as described above, and the surface function was used to identify individual nuclei, using a surface detail of 1 µm and a background subtraction of 4 µm. The split touching objects function was set at 4 µm and used to isolate individual nuclei. Individual nuclei were then tracked across time following an autoregressive motion algorithm, with a max distance of 8 µm and gap size of 0. Tracks were then manually checked for accuracy and corrected as necessary. The track function works best for early stages (below than 36 hpf) where diastolic movements are slower.

### Strain inference from the myocardial nuclei trajectories

Nuclei coordinates from nuclei track data were used to reconstruct surfaces of the heart at each stage of the cardiac cycle. For 48 hpf wild type and *amhc* embryos, systole, the state in which the myocardium is maximally contracted, was used as the elastic reference configuration for the calculation of strain. For earlier stages and the *tbx5a* morphant, the ventricle beats in a peristaltic-like motion (*115*) and hence there is no state in which the entire ventricle is maximally contracted. To determine the reference configuration, nuclear coordinates across the whole cardiac cycle were superimposed, and points that lie closest to the central axis of the point cloud were taken. Smooth and differentiable reference surfaces were constructed using the screened Poisson surface reconstruction algorithm (*116*) in MeshLab (*117*)

The displacement of nuclei at each stage of the cardiac cycle was then computed by determining the positional differences between their current position and their position in the reference state (**Fig. S7B**). A displacement field was then calculated over the full reference surface by interpolating the computed displacement data, using a spherical Gaussian interpolation facilitated by VTK (*118*) and vedo (*119*). This transforms the original discrete displacement field into a dense and continuous displacement field that accurately represents the motion of the whole myocardium over the cardiac cycle.

As nuclei are sparsely distributed, the displacement field is further smoothened to eliminate noise that can affect subsequent numerical computations, by penalizing gradients in the deformation maps. This effectively minimizes abrupt changes or discontinuities in the displacement vectors, producing a physically realistic field that captures the biomechanical behavior of the myocardium while being smooth enough for numerical computations.

To quantify how much the myocardium has stretched with respect to the elastic reference, we compute true strain, following:

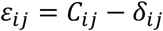

Where_*C_ij_* is the right-Cauchy deformation tensor (see the supplementary modelling section) and *δ_ij_* is the Kronecker delta (*120*).

### Average fractures map

For an average representation of the distribution of simulated cECM fractures, average fracture maps were computed from three different hearts at each stage (**Fig S8A, Fig S8E**). For this, the simulated fractures for each heart were first projected onto a two-dimensional surface. Due to the natural variability in heart size, each heart was rescaled by fitting to a common reference shape. This allows different hearts to be superimposed, whilst preserving the relative spatial distribution of fractures. Once rescaled, the projected maps were then filtered using a sigmoid function to binarize the data, classifying each region as either fractured or not, based on a predefined threshold. This ensures a consistent interpretation of fractures across each sample. To obtain the average fracture map, the binarized fracture patterns from all three independent simulations were pooled, and their mean was calculated. This accurately captures the geometric distribution of fractures whilst minimizing noise from individual variations in the simulations.

### Modelling

To investigate whether the cECM undergoes mechanical fracturing due to cyclic myocardial contractions, we developed a mathematical model treating the cECM as a thin viscoelastic sheet subjected to progressive damage. The cECM is modelled as an incompressible neo-Hookean material with an additional Newtonian viscous stress. Myocardial deformation induces deformation in the cECM, which can slide relative to the myocardium but with a frictional penalty. Damage is introduced via a scalar phase-field variable α(X,t), where α=1 represents an undamaged state and α=0 denotes complete failure. The system dynamics are governed by the following coupled equations:

(1) **Balance of forces and shape deformation.** On the deformed cECM surface,

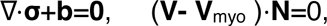

where **V** is the 3D velocity of the cECM, **σ**=g(α)**σ**_el_+2μα**d** is the Cauchy stress tensor, **σ**_el_ the elastic stress from a neoHookean material, g(α) defining material softening due to damage, **d** the strain rate, and μ the viscosity of the undamaged cECM. ∇ stands for the covariant derivative on the surface. The term **b**=−ηα**P**(**V-V**^myo^) accounts for the frictional force between the cECM and the myocardium, with η the friction coefficient, **P** the projector operator onto the tangent plane of the surface and **V**^myo^ the velocity of the myocardium. The first equation is a statement of balance of forces tangent to the cECM surface, and the second equation imposes the normal velocity of the cECM to coincide with that of the myocardium.

(2) **Time-evolution of the damage field.**

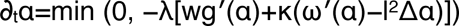

Where λ controls the damage dissipation rate, w is the elastic energy density, κ characterizes the energy released per unit area for the damage of an unstretched material, and l determines the characteristic fracture length scale.

For numerical implementation, the model equations are discretized using a finite element approach on a triangular mesh. Time evolution follows a discrete Onsager variational principle, and constraints such as myocardial movement and damage irreversibility are enforced using penalty functions. The numerical solution is computed using the *hiperlife* library (*121*). See more details in the supplementary material. For each simulation at 30, 36, 42, and 48 hours, we assume that the cECM is initially undamaged and simulate only the localization and geometry of newly formed fractures. The reference configuration for each simulation is taken as the systolic or the reconstructed maximally contracted shape at the corresponding developmental stage. This choice reflects the assumption that, over long timescales, the cECM undergoes plastic adaptation through creep, gradually adjusting its reference configuration. This is supported by our observation that the cECM retains the heart’s resting shape across different developmental stages (**Fig. S7D**) and is consistent with previous studies on the cECM of chick embryos (*61*)

Code is available at https://git.embl.de/grp-torres-sanchez/cECM-fractures. All simulations were run in the EMBL-HD HPC cluster (*122*)

### Chicken egg incubation, embryo staging and imaging

Roslin Green Eggs obtained from the National Avian Research Facility at the University of Edinburgh were incubated for 36/43/46 hours at 38°C to generate Hamburger Hamilton stage (HH) 10/11/12embryos with 10/13/16 somites (*123*). Embryos were dissected from the eggs at desired stages and washed with PBS. To image the beating heart, embryos were directly imaged post dissection with a Leica SMZ18 microscope. For confocal imaging, embryos were mounted in 35 mm diameter glass-bottomed dishes, in 1.2% (w/v) low-melting agarose / PBS containing 2 µg/µl tricaine to stop the hearts. Once the agarose was set, the dish was then filled with PBS containing 4 µg/µl tricaine. All stopped-heart images were acquired using a Zeiss LSM980 Axio Examiner confocal microscope with a Zeiss W Plan-Apochromat 10x / 1.0 DIC M27 water immersion dipping lens.

### Chicken egg drug treatments

Eggs were opened at HH10 and 7 ml of albumin was removed. 4 ml of 400 µM Nifedipine or 1% DMSO (vehicle control) was added to each egg, and eggs were closed using sticky tape. Eggs were then incubated at 38°C for 10 hours post-treatment, before being imaged on the Zeiss LSM980 as described above. This treatment is sufficient to stop the heart, and Nifedipine treated embryos at HH12 do not have beating hearts.

### Statistics

No statistical methods were used to predetermine sample size. Functional experiments were performed independently at least three times. Data used for characterisation, including chick data and data for **Fig. S7E** were performed independently at least twice. Whenever possible, blinding was performed in data collection and analysis. GraphPad Prism (v.9) was used to perform all statistical analyses and plot most graphs. Some graphs were plotted using custom Matlab or R scripts, namely the density heatmaps, strain maps and 2D cECM thickness graphs.

## Tables

**Table S1:** List of transgenic lines, cellular structures and processes labelled by them and corresponding abbreviation.

| Line | Cellular structures and processes | Abbreviation |
| --- | --- | --- |
| *Tg(myl7:BFP-CAAX)<sup>bns193</sup>* | Myocardium Membrane | *myl7:BFP-CAAX* (124) |
| *Tg(ubb:SEC-Rno.Ncan-EGFP)<sup>uq25bh</sup>* | Hyaluronan | *ubb:HA-GFP* (44) |
| *Tg(-0.8myl7:H2BmScarlet)<sup>bns534</sup>* | Myocardium Nuclei | *myl7:H2B-mScarlet* (125) |
| *Tg(gata1a:DsRed)<sup>sd2</sup>* | Blood Cells | *gata1:DsRed* (126) |
| *Tg(kdrl:NLS-mCherry)<sup>is4</sup>* | Endocardium Nuclei | *kdrl:NLS-mCherry* (127) |
| *Tg(kdrl:Hsa.HRAS-mCherry)<sup>s896</sup>* | Endocardium Membrane | *kdrl:HRAS-mCherry* (128) |
| *Tg(kdrl:eGFP)<sup>s843</sup>* | Endocardium Cytoplasm | *kdrl:eGFP* (129) |
| *Tg(myl7:KTR-Clover-H2B-mScarlet)<sup>bns565</sup>* | ERK reporter | *myl7:KTR-Clover* (130) |
| *Tg(-0.8myl7:H2B-mApple)<sup>fci604</sup>* | Myocardium Nuclei | *myl7:H2B-mApple* |
| *Tg(kdrl:LIFEACT-eGFP)<sup>s982</sup>* | Endocardium Actin | *kdrl:lifeact-GFP* (131) |
| *haf<sup>sk24</sup>* | half-hearted mutant | NA (57) |
| *nrg2a<sup>mn0237Gt</sup>* | *nrg2a* genetrap | NA (132) |

**Table S2:**
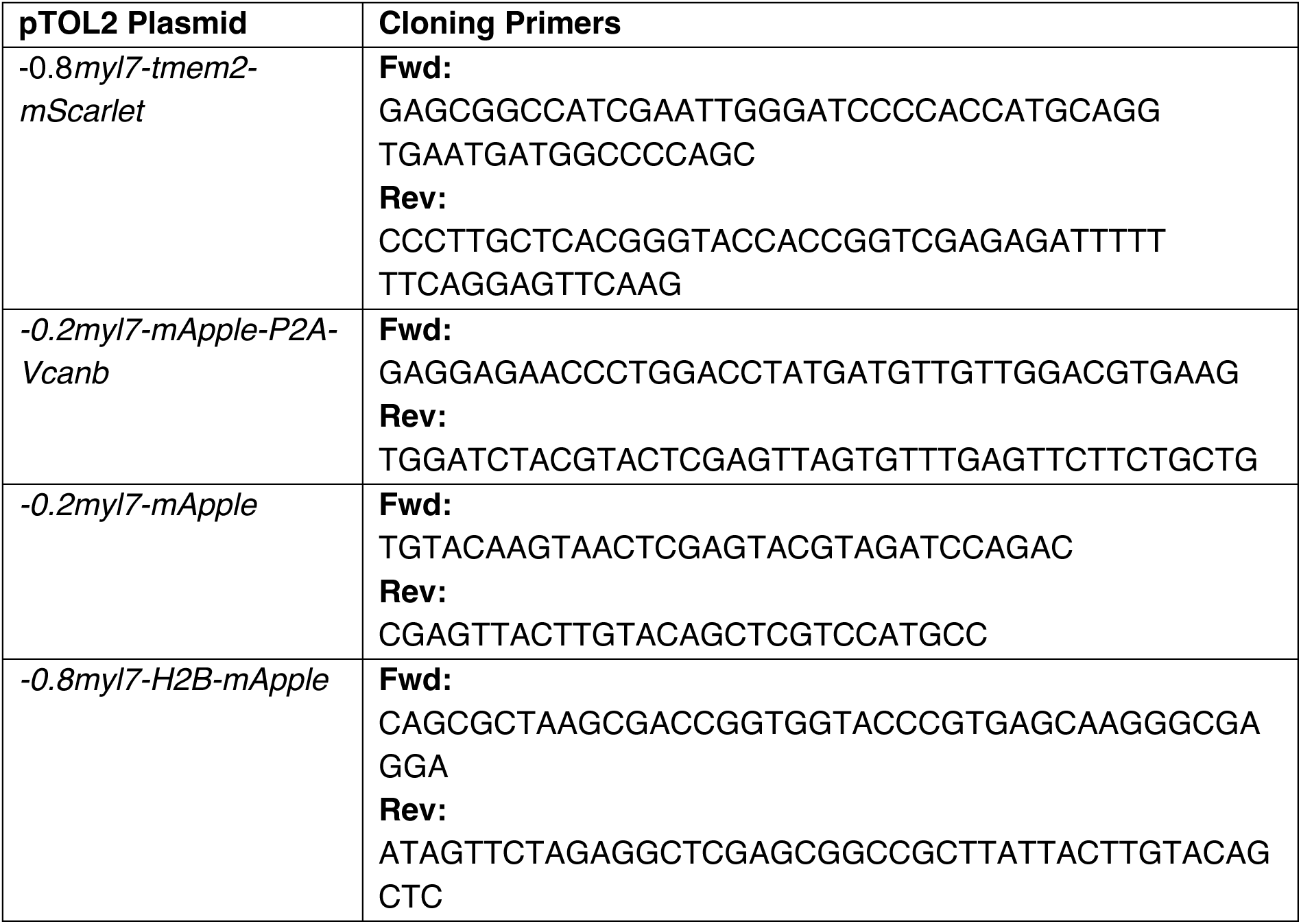
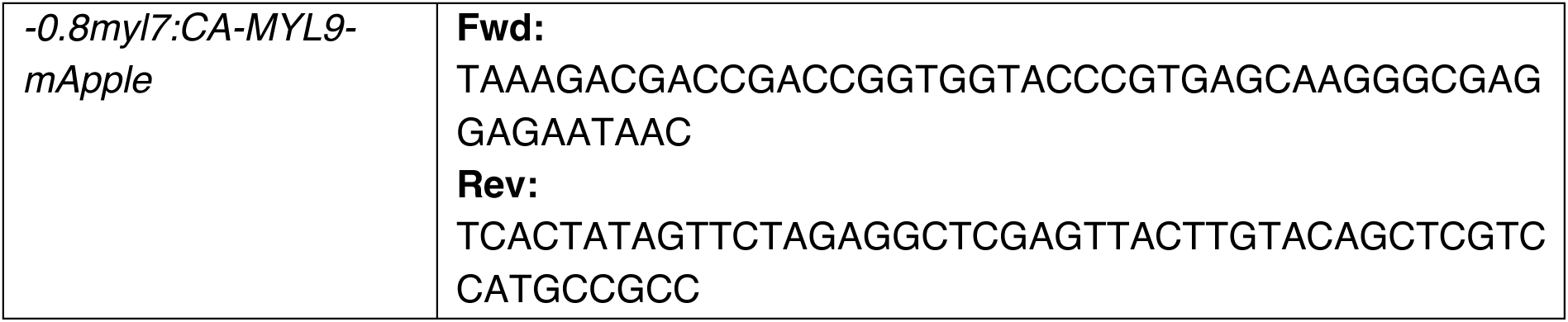
Primers used for gene cloning using In-Fusion^®^ technology.

**Table S3:**
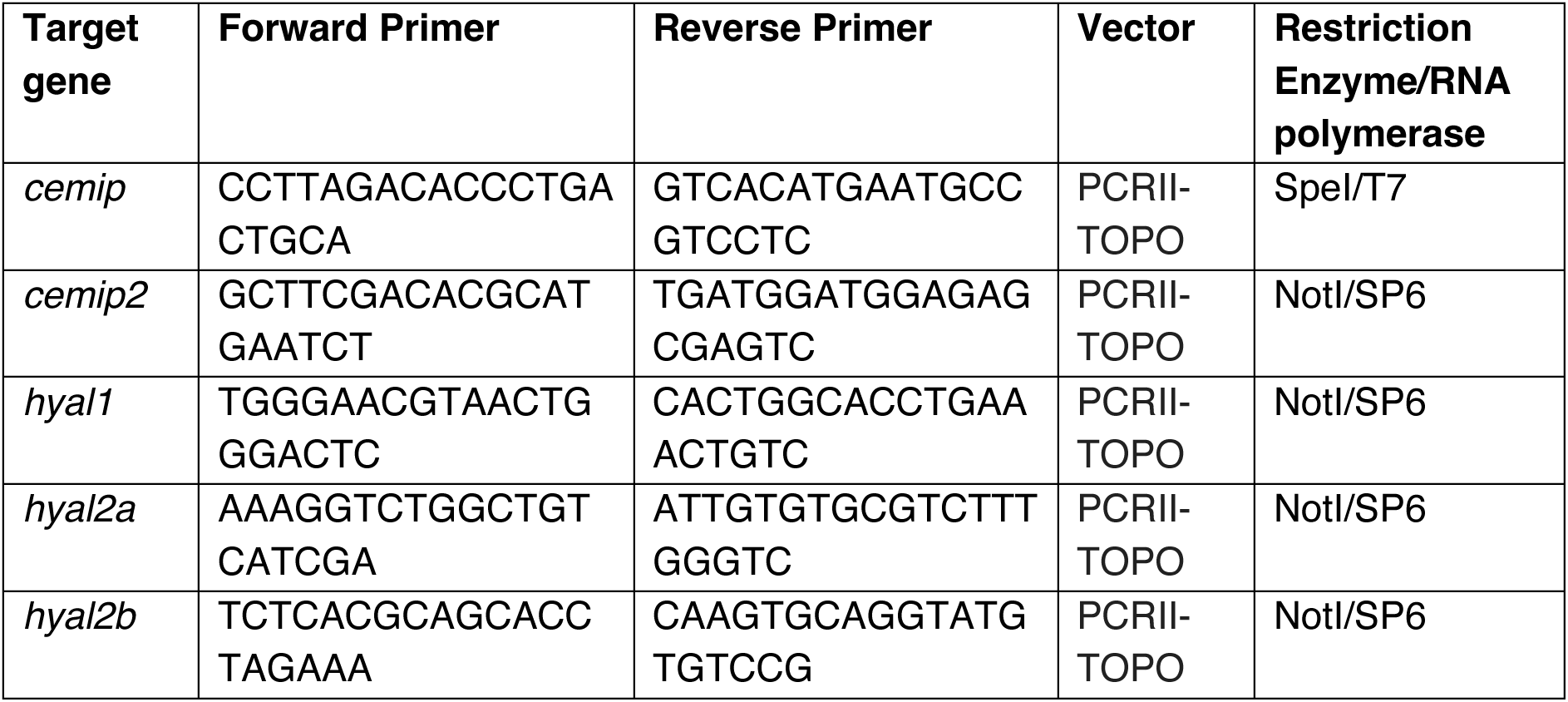
Primers used for *in situ* hybridisations.

## Reference

1. J. Briscoe, S. Small, Morphogen rules: design principles of gradient-mediated embryo patterning. Development 142, 3996–4009 (2015).

2. D. Gilmour, M. Rembold, M. Leptin, From morphogen to morphogenesis and back. Nature 541, 311–320 (2017).

3. F. Schweisguth, F. Corson, Self-Organization in Pattern Formation. Developmental Cell 49, 659–677 (2019).

4. T. G. R. Andrews, R. Priya, The Mechanics of Building Functional Organs. Cold Spring Harb Perspect Biol, (2024).

5. C. Collinet, T. Lecuit, Programmed and self-organized flow of information during morphogenesis. Nature Reviews Molecular Cell Biology 22, 245–265 (2021).

6. C. J. Chan, C.-P. Heisenberg, T. Hiiragi, Coordination of Morphogenesis and Cell-Fate Specification in Development. Current Biology 27, R1024–R1035 (2017).

7. I. Martyn, Z. J. Gartner, Expanding the boundaries of synthetic development. Dev Biol 474, 62–70 (2021).

8. J. Li et al., The Strength of Mechanical Forces Determines the Differentiation of Alveolar Epithelial Cells. Dev Cell 44, 297–312.e295 (2018).

9. S. B. Dahl-Jensen et al., Deconstructing the principles of ductal network formation in the pancreas. PLoS Biol 16, e2002842 (2018).

10. H. Fukui et al., Bioelectric signaling and the control of cardiac cell identity in response to mechanical forces. Science 374, 351–354 (2021).

11. F. Tessadori et al., Twisting of the zebrafish heart tube during cardiac looping is a tbx5-dependent and tissue-intrinsic process. eLife 10, e61733 (2021).

12. D. Staudt, D. Stainier, Uncovering the Molecular and Cellular Mechanisms of Heart Development Using the Zebrafish. Annual Review of Genetics 46, 397–418 (2012).

13. F. Gunawan, R. Priya, D. Y. R. Stainier, Sculpting the heart: Cellular mechanisms shaping valves and trabeculae. Curr Opin Cell Biol 73, 26–34 (2021).

14. V. A. Lombardo et al., Morphogenetic control of zebrafish cardiac looping by Bmp signaling. Development 146, (2019).

15. E. S. Noël et al., A Nodal-independent and tissue-intrinsic mechanism controls heart-looping chirality. Nature Communications 4, 2754 (2013).

16. J. Liu et al., A dual role for ErbB2 signaling in cardiac trabeculation. Development 137, 3867–3875 (2010).

17. R. Priya et al., Tension heterogeneity directs form and fate to pattern the myocardial wall. Nature 588, 130–134 (2020).

18. G. Del Monte-Nieto et al., Control of cardiac jelly dynamics by NOTCH1 and NRG1 defines the building plan for trabeculation. Nature 557, 439–445 (2018).

19. L. A. Samsa, B. Yang, J. Liu, Embryonic cardiac chamber maturation: Trabeculation, conduction, and cardiomyocyte proliferation. Am J Med Genet C Semin Med Genet 163c, 157–168 (2013).

20. S. J. Rasouli, D. Y. R. Stainier, Regulation of cardiomyocyte behavior in zebrafish trabeculation by Neuregulin 2a signaling. Nat Commun 8, 15281 (2017).

21. J. Qi et al., Apelin signaling dependent endocardial protrusions promote cardiac trabeculation in zebrafish. Elife 11, (2022).

22. A. Sizarov et al., Formation of the building plan of the human heart: morphogenesis, growth, and differentiation. Circulation 123, 1125–1135 (2011).

23. A. F. M. Moorman, V. M. Christoffels, Cardiac Chamber Formation: Development, Genes, and Evolution. Physiological Reviews 83, 1223–1267 (2003).

24. T. G. Andrews et al., Multiscale mechanics drive functional maturation of the vertebrate heart. bioRxiv, 2024.2007.2024.604962 (2024).

25. G. Captur et al., Morphogenesis of myocardial trabeculae in the mouse embryo. J Anat 229, 314–325 (2016).

26. G. del Monte-Nieto, J. W. Fischer, D. J. Gorski, R. P. Harvey, J. C. Kovacic, Basic Biology of Extracellular Matrix in the Cardiovascular System, Part 1/4: JACC Focus Seminar. Journal of the American College of Cardiology 75, 2169-2188 (2020).

27. V. Jiménez-Amilburu et al., In Vivo Visualization of Cardiomyocyte Apicobasal Polarity Reveals Epithelial to Mesenchymal-like Transition during Cardiac Trabeculation. Cell Rep 17, 2687–2699 (2016).

28. D. W. Staudt et al., High-resolution imaging of cardiomyocyte behavior reveals two distinct steps in ventricular trabeculation. Development 141, 585–593 (2014).

29. V. Uribe et al., In vivo analysis of cardiomyocyte proliferation during trabeculation. Development 145, (2018).

30. A. F. Moorman, V. M. Christoffels, Cardiac chamber formation: development, genes, and evolution. Physiol Rev 83, 1223–1267 (2003).

31. H. M. T. Choi et al., Third-generation in situ hybridization chain reaction: multiplexed, quantitative, sensitive, versatile, robust. Development 145, (2018).

32. J. Grego-Bessa et al., Nrg1 Regulates Cardiomyocyte Migration and Cell Cycle in Ventricular Development. Circ Res 133, 927–943 (2023).

33. C. de la Cova, R. Townley, S. Regot, I. Greenwald, A Real-Time Biosensor for ERK Activity Reveals Signaling Dynamics during C. elegans Cell Fate Specification. Dev Cell 42, 542–553 e544 (2017).

34. C. J. Derrick, E. S. Noël, The ECM as a driver of heart development and repair. Development 148, (2021).

35. A. Gentile et al., Mechanical forces remodel the cardiac extracellular matrix during zebrafish development. Development 151, (2024).

36. N. Khalilgharibi, Y. Mao, To form and function: on the role of basement membrane mechanics in tissue development, homeostasis and disease. Open Biology 11, 200360 (2021).

37. M. D. Diaz-de-la-Loza, B. M. Stramer, The extracellular matrix in tissue morphogenesis: No longer a backseat driver. Cells Dev 177, 203883 (2024).

38. R. Sekiguchi, K. M. Yamada, Basement Membranes in Development and Disease. Curr Top Dev Biol 130, 143–191 (2018).

39. E. Steed et al., klf2a couples mechanotransduction and zebrafish valve morphogenesis through fibronectin synthesis. Nature Communications 7, 11646 (2016).

40. E. J. G. Pollitt, J. Sánchez-Posada, C. M. Snashall, C. J. Derrick, E. S. Noël, Llgl1 mediates timely epicardial emergence and establishment of an apical laminin sheath around the trabeculating cardiac ventricle. Development 151, (2024).

41. C. J. Derrick et al., Asymmetric Hapln1a drives regionalized cardiac ECM expansion and promotes heart morphogenesis in zebrafish development. Cardiovasc Res 118, 226–240 (2022).

42. A. Nakamura, F. J. Manasek, Experimental studies of the shape and structure of isolated cardiac jelly. J Embryol Exp Morphol 43, 167–183 (1978).

43. A. Nakamura, F. J. Manasek, An experimental study of the relation of cardiac jelly to the shape of the early chick embryonic heart. Development 65, 235–256 (1981).

44. D. R. Grassini et al., Nppa and Nppb act redundantly during zebrafish cardiac development to confine AVC marker expression and reduce cardiac jelly volume. Development 145, (2018).

45. J. E. De Angelis et al., Tmem2 Regulates Embryonic Vegf Signaling by Controlling Hyaluronic Acid Turnover. Dev Cell 40, 123–136 (2017).

46. J. Sanchez-Posada, C. J. Derrick, E. S. Noel, morphoHeart: A quantitative tool for integrated 3D morphometric analyses of heart and ECM during embryonic development. PLoS Biol 23, e3002995 (2025).

47. C. Frantz, K. M. Stewart, V. M. Weaver, The extracellular matrix at a glance. J Cell Sci 123, 4195–4200 (2010).

48. Y. Mori, S. Smith, J. Wang, A. Munjal, Versican controlled by Lmx1b regulates hyaluronate density and hydration for semicircular canal morphogenesis. bioRxiv, 2024.2005.2007.592968 (2024).

49. T. D. Camenisch, J. A. Schroeder, J. Bradley, S. E. Klewer, J. A. McDonald, Heart-valve mesenchyme formation is dependent on hyaluronan-augmented activation of ErbB2–ErbB3 receptors. Nature Medicine 8, 850–855 (2002).

50. T. D. Camenisch et al., Disruption of hyaluronan synthase-2 abrogates normal cardiac morphogenesis and hyaluronan-mediated transformation of epithelium to mesenchyme. J Clin Invest 106, 349–360 (2000).

51. H. Yamamoto et al., A mammalian homolog of the zebrafish transmembrane protein 2 (TMEM2) is the long-sought-after cell-surface hyaluronidase. J Biol Chem 292, 7304–7313 (2017).

52. T. Narita et al., TMEM2 is a bona fide hyaluronidase possessing intrinsic catalytic activity. J Biol Chem 299, 105120 (2023).

53. R. Totong et al., The novel transmembrane protein Tmem2 is essential for coordination of myocardial and endocardial morphogenesis. Development 138, 4199–4205 (2011).

54. J. Bakkers et al., Has2 is required upstream of Rac1 to govern dorsal migration of lateral cells during zebrafish gastrulation. Development 131, 525–537 (2004).

55. T. L. Anderson, Fracture Mechanics: Fundamentals and Applications, Fourth Edition (4th ed.). CRC Press, (2017).

56. J. Gierten et al., Automated high-throughput heartbeat quantification in medaka and zebrafish embryos under physiological conditions. Sci Rep 10, 2046 (2020).

57. H. J. Auman et al., Functional Modulation of Cardiac Form through Regionally Confined Cell Shape Changes. PLOS Biology 5, e53 (2007).

58. P. Sidhwani, D. Yelon, Fluid forces shape the embryonic heart: Insights from zebrafish. Curr Top Dev Biol 132, 395–416 (2019).

59. J. Vermot et al., Reversing blood flows act through klf2a to ensure normal valvulogenesis in the developing heart. PLoS Biol 7, e1000246 (2009).

60. N. Ebrahimi et al., A method for investigating spatiotemporal growth patterns at cell and tissue levels during C-looping in the embryonic chick heart. iScience 25, (2022).

61. J. Yao et al., Viscoelastic material properties of the myocardium and cardiac jelly in the looping chick heart. J Biomech Eng 134, 024502 (2012).

62. E. Berdougo, H. Coleman, D. H. Lee, D. Y. Stainier, D. Yelon, Mutation of weak atrium/atrial myosin heavy chain disrupts atrial function and influences ventricular morphogenesis in zebrafish. Development 130, 6121–6129 (2003).

63. C. M. Nelson, Geometric control of tissue morphogenesis. Biochimica et Biophysica Acta (BBA) - Molecular Cell Research 1793, 903–910 (2009).

64. C. M. Nelson et al., Emergent patterns of growth controlled by multicellular form and mechanics. Proc Natl Acad Sci U S A 102, 11594–11599 (2005).

65. V. D. Varner, J. P. Gleghorn, E. Miller, D. C. Radisky, C. M. Nelson, Mechanically patterning the embryonic airway epithelium. Proceedings of the National Academy of Sciences 112, 9230–9235 (2015).

66. C. M. Nelson, M. M. Vanduijn, J. L. Inman, D. A. Fletcher, M. J. Bissell, Tissue geometry determines sites of mammary branching morphogenesis in organotypic cultures. Science 314, 298–300 (2006).

67. E. W. Gehrels, B. Chakrabortty, M.-E. Perrin, M. Merkel, T. Lecuit, Curvature gradient drives polarized tissue flow in the *Drosophila* embryo. Proceedings of the National Academy of Sciences 120, e2214205120 (2023).

68. A. Villedieu et al., Homeotic compartment curvature and tension control spatiotemporal folding dynamics. Nature Communications 14, 594 (2023).

69. T. Hirashima, M. Matsuda, ERK-mediated curvature feedback regulates branching morphogenesis in lung epithelial tissue. Curr Biol 34, 683–696.e686 (2024).

70. N. E. Vlahakis, R. D. Hubmayr, Response of alveolar cells to mechanical stress. Curr Opin Crit Care 9, 2–8 (2003).

71. K. Uenishi, Fracture dynamics of solid materials: from particles to the globe. Philosophical Transactions of the Royal Society A: Mathematical, Physical and Engineering Sciences 379, 20200122 (2021).

72. K. Ravi-Chandar, in Comprehensive Structural Integrity (Second Edition), M. H. F. Aliabadi, W. O. Soboyejo, Eds. (Elsevier, Oxford, 2003), pp. 117-192.

73. G. N. Santos-Durán, R. L. Cooper, E. Jahanbakhsh, G. Timin, M. C. Milinkovitch, Self-organized patterning of crocodile head scales by compressive folding. Nature 637, 375–383 (2025).

74. M. C. Milinkovitch et al., Crocodile Head Scales Are Not Developmental Units But Emerge from Physical Cracking. Science 339, 78–81 (2013).

75. J. G. Dumortier et al., Hydraulic fracturing and active coarsening position the lumen of the mouse blastocyst. Science 365, 465–468 (2019).

76. J. Duque et al., Rupture strength of living cell monolayers. Nature Materials 23, 1563–1574 (2024).

77. A. Bonfanti, J. Duque, A. Kabla, G. Charras, Fracture in living tissues. Trends Cell Biol 32, 537–551 (2022).

78. E. W. Levy, I. Leite, B. W. Joyce, S. Y. Shvartsman, E. Posfai, A tug-of-war between germ cell motility and intercellular bridges controls germline cyst formation in mice. Current Biology, (2024).

79. V. N. Prakash, M. S. Bull, M. Prakash, Motility-induced fracture reveals a ductile-to-brittle crossover in a simple animal’s epithelia. Nature Physics 17, 504–511 (2021).

80. W. Xi, T. B. Saw, D. Delacour, C. T. Lim, B. Ladoux, Material approaches to active tissue mechanics. Nature Reviews Materials 4, 23–44 (2019).

81. D. A. C. Walma, K. M. Yamada, The extracellular matrix in development. Development 147, (2020).

82. K. H. Palmquist et al., Reciprocal cell-ECM dynamics generate supracellular fluidity underlying spontaneous follicle patterning. Cell 185, 1960–1973.e1911 (2022).

83. C. Kyprianou et al., Basement membrane remodelling regulates mouse embryogenesis. Nature 582, 253–258 (2020).

84. V. F. Fiore et al., Mechanics of a multilayer epithelium instruct tumour architecture and function. Nature 585, 433–439 (2020).

85. N. Bansaccal et al., The extracellular matrix dictates regional competence for tumour initiation. Nature, (2023).

86. K. M. Wisdom et al., Matrix mechanical plasticity regulates cancer cell migration through confining microenvironments. Nature Communications 9, 4144 (2018).

87. A. Glentis et al., Cancer-associated fibroblasts induce metalloprotease-independent cancer cell invasion of the basement membrane. Nature Communications 8, 924 (2017).

88. J. S. Harunaga, A. D. Doyle, K. M. Yamada, Local and global dynamics of the basement membrane during branching morphogenesis require protease activity and actomyosin contractility. Dev Biol 394, 197–205 (2014).

89. J. Chang et al., Cell volume expansion and local contractility drive collective invasion of the basement membrane in breast cancer. Nature Materials 23, 711–722 (2024).

90. A. Munjal, E. Hannezo, T. Y. Tsai, T. J. Mitchison, S. G. Megason, Extracellular hyaluronate pressure shaped by cellular tethers drives tissue morphogenesis. Cell 184, 6313–6325.e6318 (2021).

91. H. Hamada, Hyaluronan Works on the Right for Directional Gut Looping. Developmental Cell 46, 525–526 (2018).

92. B. D. Sanketi et al., Pitx2 patterns an accelerator-brake mechanical feedback through latent TGFβ to rotate the gut. Science 377, eabl3921 (2022).

93. H. Vignes et al., Extracellular mechanical forces drive endocardial cell volume decrease during zebrafish cardiac valve morphogenesis. Developmental Cell 57, 598–609.e595 (2022).

94. M. Chugh, A. Munjal, S. G. Megason, Hydrostatic pressure as a driver of cell and tissue morphogenesis. Semin Cell Dev Biol 131, 134–145 (2022).

95. A. P. Spicer, J. Y. L. Tien, Hyaluronan and morphogenesis. Birth Defects Research Part C: Embryo Today: Reviews 72, 89–108 (2004).

96. B. P. Toole, Hyaluronan in morphogenesis. Seminars in Cell & Developmental Biology 12, 79–87 (2001).

97. H. A. Messal et al., Tissue curvature and apicobasal mechanical tension imbalance instruct cancer morphogenesis. Nature 566, 126–130 (2019).

98. J. Folkman, H. P. Greenspan, Influence of geometry on control of cell growth. Biochim Biophys Acta 417, 211–236 (1975).

99. N. Gjorevski et al., Tissue geometry drives deterministic organoid patterning. Science 375, eaaw9021 (2022).

100. J. V. Veenvliet et al., Mouse embryonic stem cells self-organize into trunk-like structures with neural tube and somites. Science 370, (2020).

101. E. Karzbrun et al., Human neural tube morphogenesis in vitro by geometric constraints. Nature 599, 268–272 (2021).

102. K. Kawakami, Tol2: a versatile gene transfer vector in vertebrates. Genome Biology 8, S7 (2007).

103. K. Kawakami, K. Asakawa, A. Muto, H. Wada, Tol2-mediated transgenesis, gene trapping, enhancer trapping, and Gal4-UAS system. Methods Cell Biol 135, 19–37 (2016).

104. L. A. Samsa et al., Cardiac contraction activates endocardial Notch signaling to modulate chamber maturation in zebrafish. Development 142, 4080–4091 (2015).

105. C. Peshkovsky, R. Totong, D. Yelon, Dependence of cardiac trabeculation on neuregulin signaling and blood flow in zebrafish. Dev Dyn 240, 446–456 (2011).

106. E. Heckel et al., Oscillatory Flow Modulates Mechanosensitive klf2a Expression through trpv4 and trpp2 during Heart Valve Development. Curr Biol 25, 1354–1361 (2015).

107. D. M. Garrity, S. Childs, M. C. Fishman, The heartstrings mutation in zebrafish causes heart/fin Tbx5 deficiency syndrome. Development 129, 4635–4645 (2002).

108. T. K. Ghosh et al., Acetylation of TBX5 by KAT2B and KAT2A regulates heart and limb development. Journal of Molecular and Cellular Cardiology 114, 185–198 (2018).

109. K. A. Smith et al., Transmembrane protein 2 (Tmem2) is required to regionally restrict atrioventricular canal boundary and endocardial cushion development. Development 138, 4193–4198 (2011).

110. J. D. Moulton, Using morpholinos to control gene expression. Curr Protoc Nucleic Acid Chem **Chapter 4**, Unit 4 30 (2007).

111. J. D. Moulton, Using Morpholinos to Control Gene Expression. Curr Protoc Nucleic Acid Chem 68, 4 30 31–34 30 29 (2017).

112. M. Liebling et al., Rapid three-dimensional imaging and analysis of the beating embryonic heart reveals functional changes during development. Developmental Dynamics 235, 2940–2948 (2006).

113. E. Biot et al., in 2008 5th IEEE International Symposium on Biomedical Imaging: From Nano to Macro. (2008), pp. 975-978.

114. K. Abu Nahia et al., scRNA-seq reveals the diversity of the developing cardiac cell lineage and molecular players in heart rhythm regulation. iScience 27, 110083 (2024).

115. M. C. Fishman, K. R. Chien, Fashioning the vertebrate heart: earliest embryonic decisions. Development 124, 2099–2117 (1997).

116. M. Kazhdan, H. Hoppe, Screened poisson surface reconstruction. ACM Trans. Graph. 32, Article 29 (2013).

117. P. Cignoni et al., MeshLab: an Open-Source Mesh Processing Tool. (2008), vol. 1, pp. 129–136.

118. W. M. Schroeder, Ken; Lorensen, Bill The Visualization Toolkit (4th ed.). (2006).

119. D. G. Musy M, Sullivan B, Vedo, a Python Module for Scientific Visualization and Analysis of 3D Objects and Point Clouds Based on Vtk, version 2021.0.2 Visualization Toolkit., (2019).

120. J. E. a. H. Marsden, Mathematical foundations of elasticity. Dover Publications, New York., (1994).

121. V. G. Santos-Oliván D, Torres-Sánchez A, hiperlife. Zenodo. (2025).

122. J. Pečar, Lueck, R., & Wahlers, M., European Molecular Biology Laboratory, (2020).

123. V. Hamburger, H. L. Hamilton, A series of normal stages in the development of the chick embryo. 1951. Dev Dyn 195, 231-272 (1992).

124. A. Guerra et al., Distinct myocardial lineages break atrial symmetry during cardiogenesis in zebrafish. eLife 7, e32833 (2018).

125. G. L. M. Boezio et al., The developing epicardium regulates cardiac chamber morphogenesis by promoting cardiomyocyte growth. Disease Models & Mechanisms 16, (2022).

126. D. Traver et al., Transplantation and in vivo imaging of multilineage engraftment in zebrafish bloodless mutants. Nature Immunology 4, 1238–1246 (2003).

127. Y. Wang et al., Moesin1 and Ve-cadherin are required in endothelial cells during in vivo tubulogenesis. Development 137, 3119–3128 (2010).

128. N. C. Chi et al., Foxn4 directly regulates tbx2b expression and atrioventricular canal formation. Genes Dev 22, 734–739 (2008).

129. L. D’Amico, I. C. Scott, B. Jungblut, D. Y. R. Stainier, A Mutation in Zebrafish *hmgcr1b* Reveals a Role for Isoprenoids in Vertebrate Heart-Tube Formation. Current Biology 17, 252-259 (2007).

130. J. Qi et al., Apelin signaling dependent endocardial protrusions promote cardiac trabeculation in zebrafish. eLife 11, e73231 (2022).

131. B. Vanhollebeke et al., Tip cell-specific requirement for an atypical Gpr124- and Reck-dependent Wnt/β-catenin pathway during brain angiogenesis. eLife 4, e06489 (2015).

132. S. E. Westcot et al., Protein-Trap Insertional Mutagenesis Uncovers New Genes Involved in Zebrafish Skin Development, Including a Neuregulin 2a-Based ErbB Signaling Pathway Required during Median Fin Fold Morphogenesis. PLoS One 10, e0130688 (2015).

