## Supplementary Figures for "Mechanical fracturing of the extracellular matrix patterns the vertebrate heart"

#### Chan et al. Supplementary Figure 1

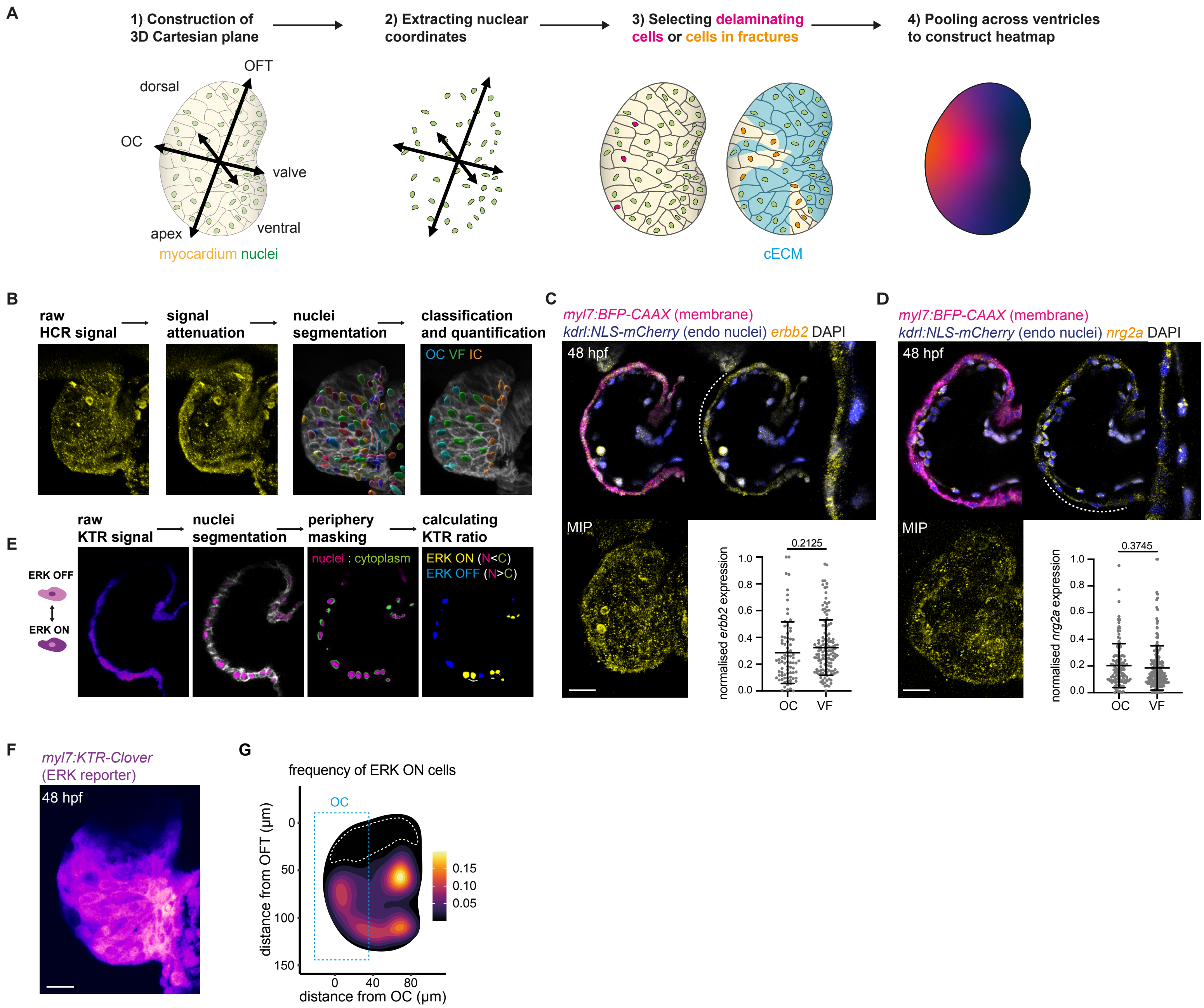

#### Chan et al. Supplementary Figure 2

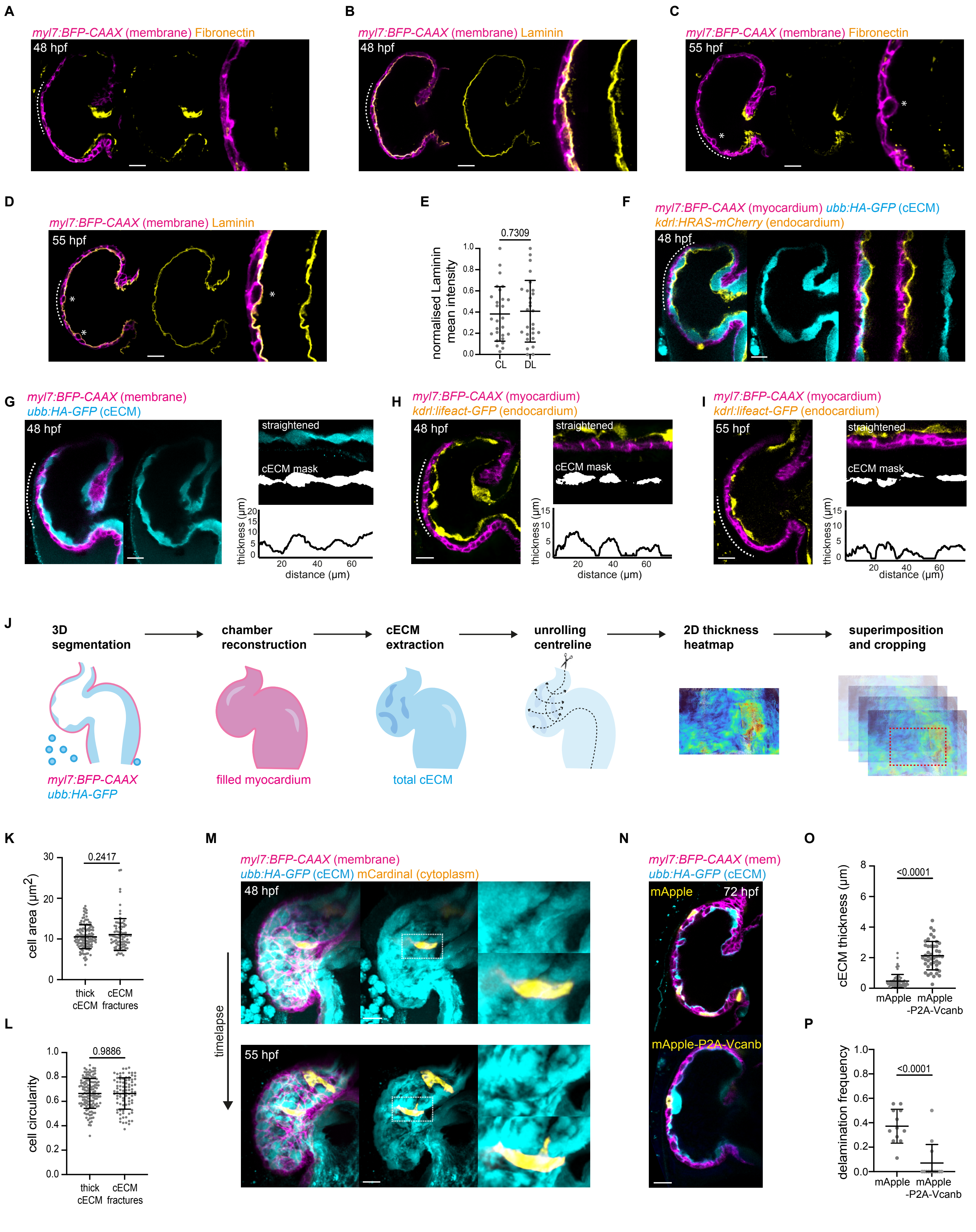

#### Chan et al. Supplementary Figure 3

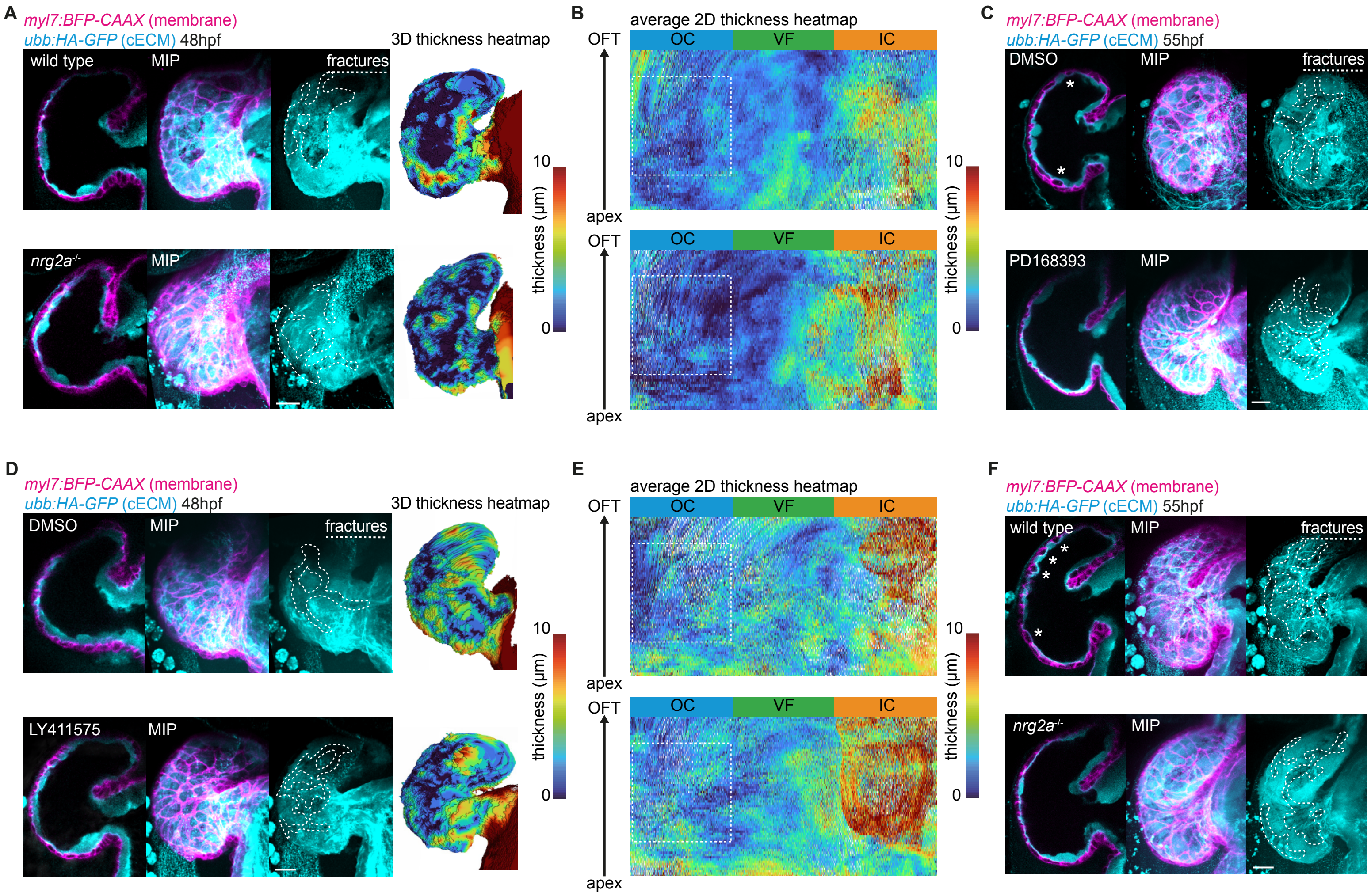

### Chan et al. Supplementary Figure 4

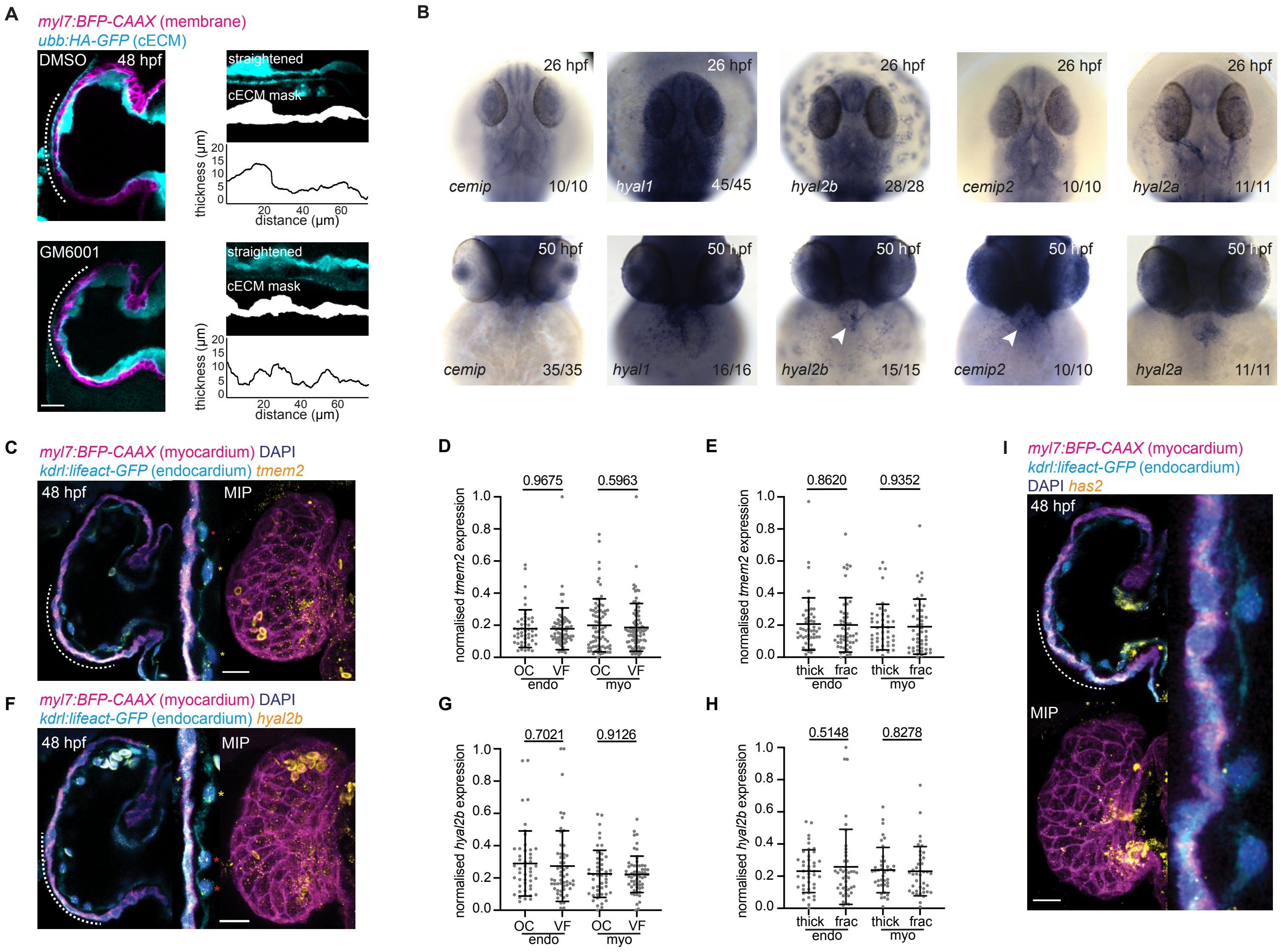

### Chan et al. Supplementary Figure 5

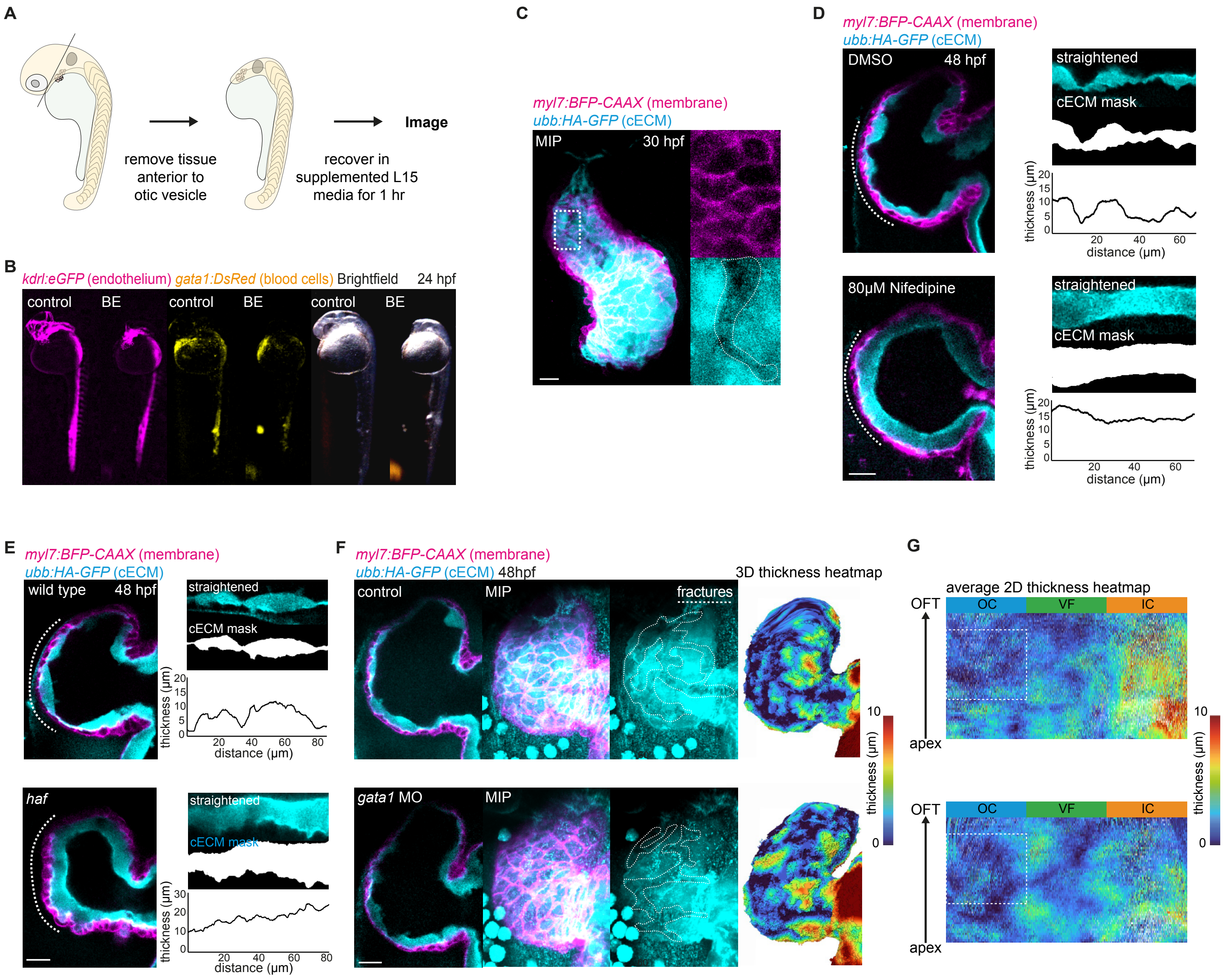

### Chan et al. Supplementary Figure 6

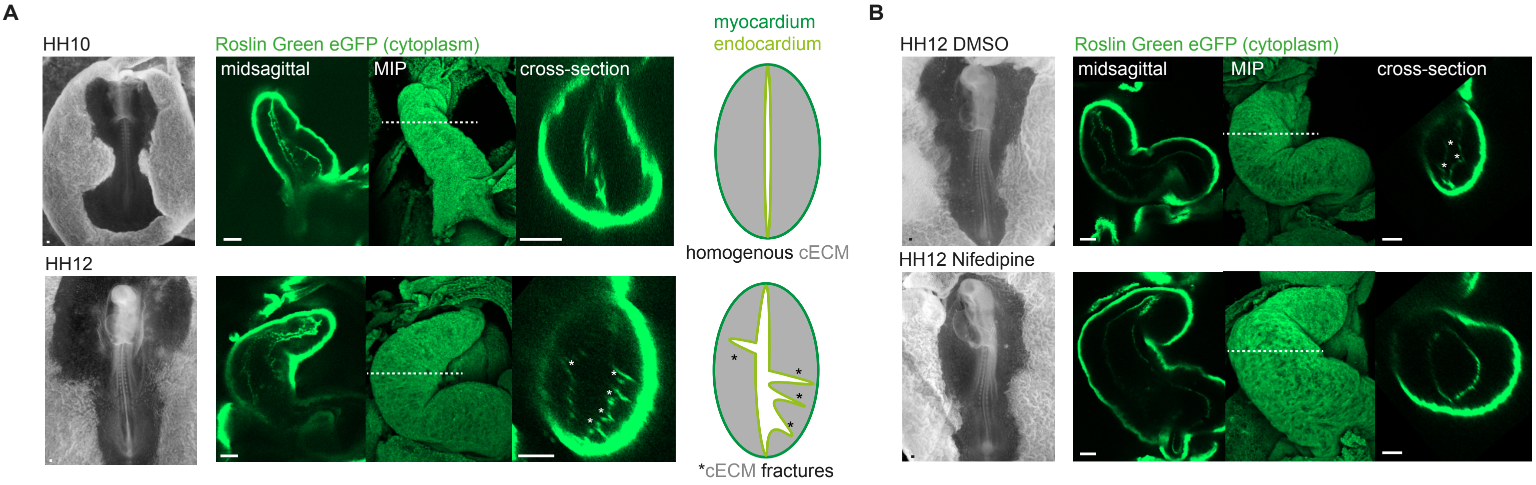

### Chan et al. Supplementary Figure 7

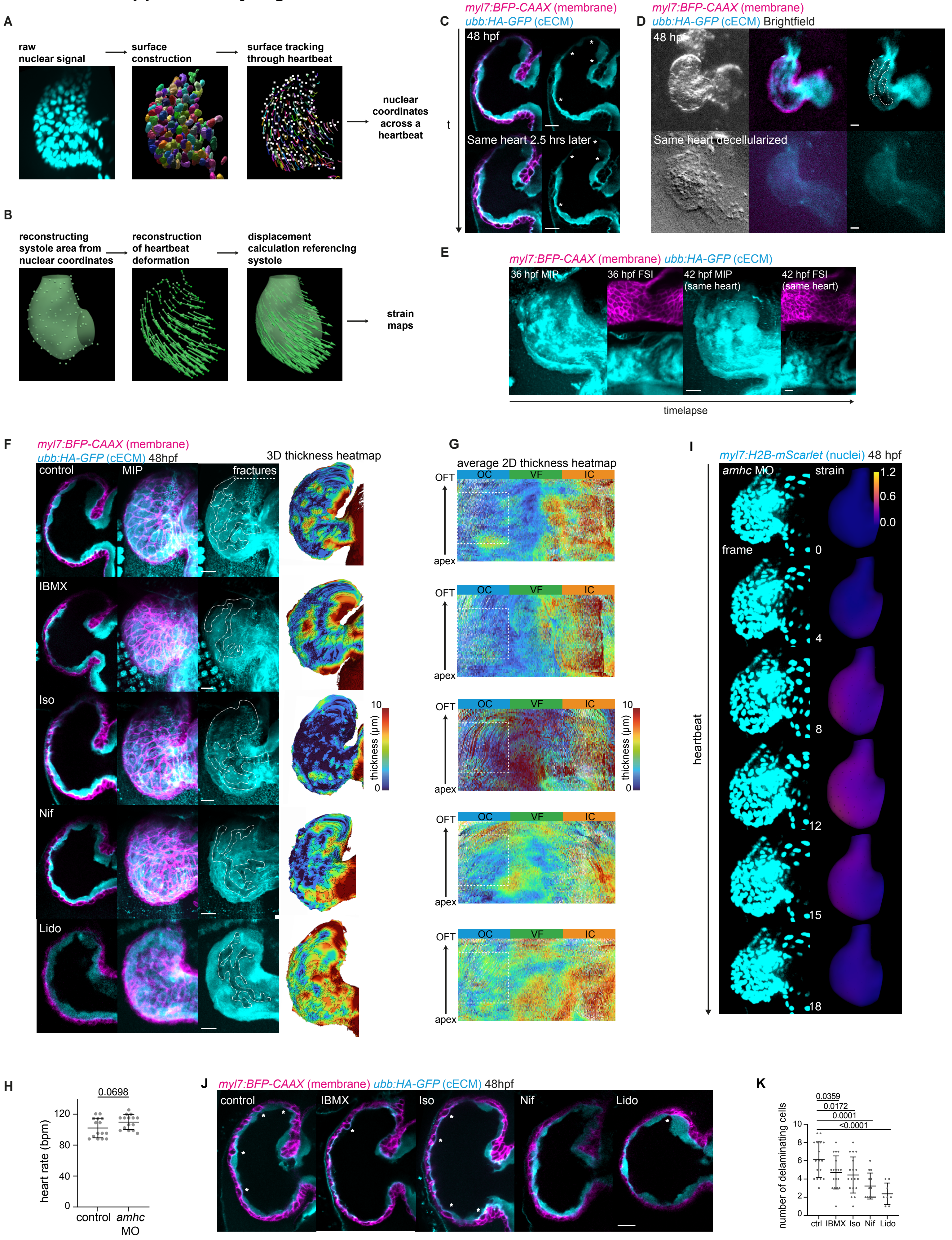

### Chan et al. Supplementary Figure 8

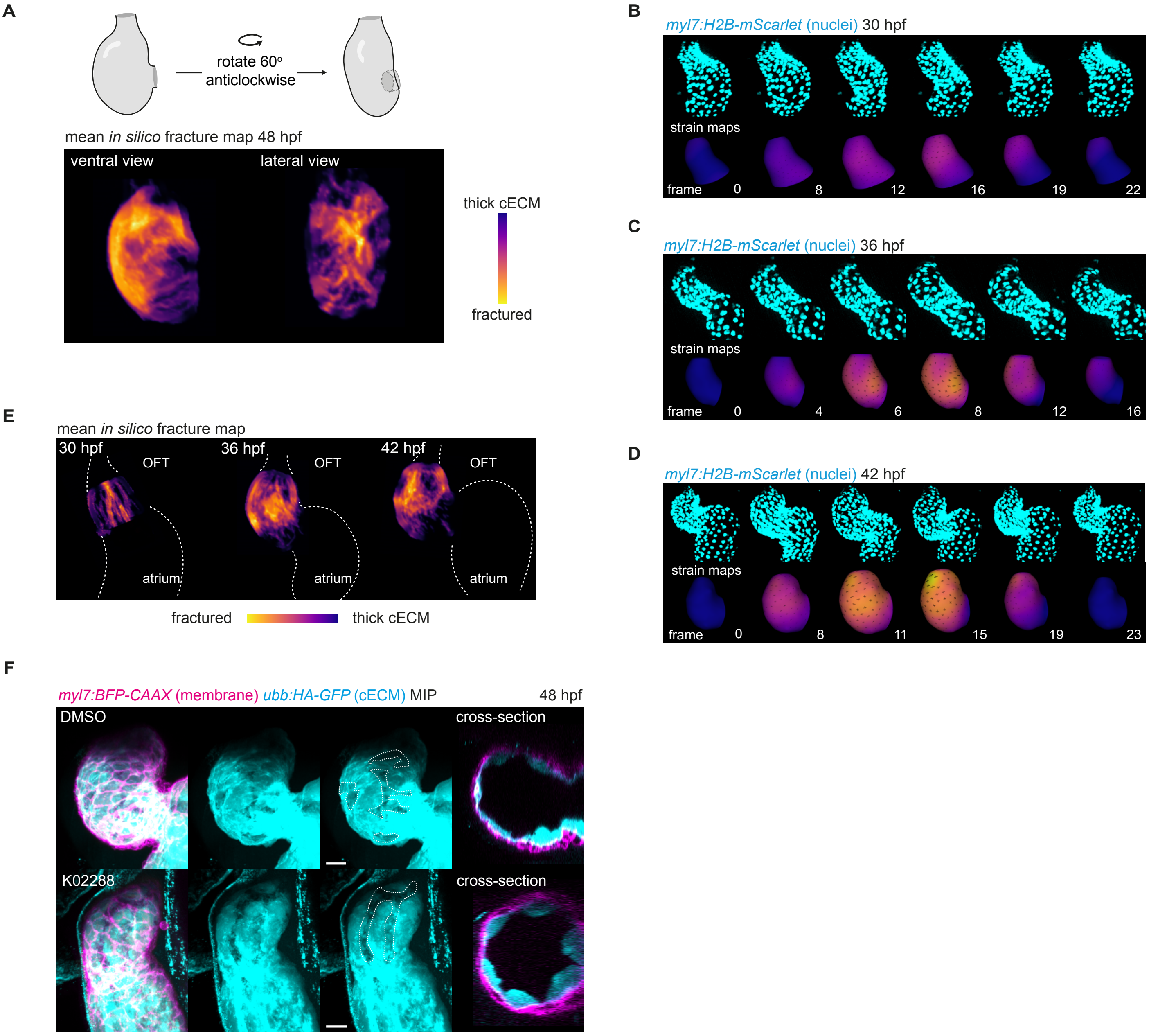
