## Supplementary material for "Mechanical fracturing of the extracellular matrix patterns the vertebrate heart": Chan et al-supplementary modelling

#### Mathematical model of the cardiac ECM and numerical implementation

To determine whether the patterns observed in the cardiac ECM (cECM) can be attributed to mechanical fracturing, we develop a mathematical model that treats the cECM as a viscoelastic thin sheet that undergoes persistent damage due to cyclic stretching. We assume that the shape of the cECM is prescribed by the deformations caused by the myocardium, and allow cECM to rearrange tangentially with respect to the myocardium by dissipating energy due to friction. Damage is accumulated over stretching cycles due to heart contractility, leading to softening and, eventually, to the emergence of fractures. To define the dynamics, we follow Onsager's variational formalism, which provides thermodynamically consistent governing equations from a variational principle [1, 2]. In the following sections, we describe the details of the mathematical formulation as well as its numerical implementation.

In our model, we do not explicitly describe the long-term creep behaviour of the cECM, which allows gradual reconfiguration in response to the evolving heart shape. Instead, we focus on the generation of damage and the emergence of fractures in small time-windows during development, where such effects can be neglected.

#### 1 Thin viscoelastic sheet without damage

To build a clear foundation for the model, we begin by formulating the behaviour of the cECM as a viscoelastic sheet in the absence of damage. This allows us to establish the fundamental mechanical framework before extending the model in the next section to incorporate the effects of damage and fracture. Because of the disparity between the thickness of the cECM ($\sim 5\,\mu\mathrm{m}$) and the size of the ventricle ($\sim 100\,\mu\mathrm{m}$) we model the cECM as a thin sheet. Its reference configuration at equilibrium is characterised by the surface $\Gamma_R$ which we parametrise with $\boldsymbol{\Phi}_\mathrm{R}(s^1, s^2)$, where $s^1$ and $s^2$ are surface coordinates. Because we do not explicitly model the long-term creep behaviour of the cECM, in our model the elastic reference configuration of the cECM is different at different developmental stages, and we take it to be its configuration at systole. At time $t$, the deformed configuration is characterised by a Lagrangian parametrisation $\boldsymbol{\Phi}(s^1, s^2, t)$ of the deformed surface $\Gamma_t$. Thus, for fixed $s^1_*$ and $s^2_*$, $\boldsymbol{\Phi}(s^1_*, s^2_*, t)$ is a curve that follows the trajectory of a material particle. The elastic energy stored due to the deformation can be written as [3]

$$\mathcal{F}[\boldsymbol{\Phi}] = \int_{\Gamma_R} w(I, J) dS, \tag{1}$$

where

$$I = \operatorname{tr} \boldsymbol{C} \text{ and } J = \sqrt{\det \boldsymbol{C}}, \tag{2}$$

are the invariants of the right-Cauchy deformation tensor $\boldsymbol{C}$, and $w(I, J)$ is the elastic energy density. In the basis $\boldsymbol{e}_{\mathrm{R}a} = \partial_b \boldsymbol{\Phi}_\mathrm{R}$ of the tangent space of $\Gamma_\mathrm{R}$ at $\boldsymbol{x}_\mathrm{R} = \boldsymbol{\Phi}_\mathrm{R}(s^1, s^2)$ the components of $\boldsymbol{C}$ are $C_{ab} = \boldsymbol{e}_a \cdot \boldsymbol{e}_b$ where $\boldsymbol{e}_a = \partial_a \boldsymbol{\Phi}$ and $\cdot$ denotes the scalar product in Euclidean space. By making $w$

a function of $I$ and $J$, we assume that the cECM behaves as an isotropic hyperelastic material. In particular, for simplicity we assume a neoHookean energy density of the form

$$w(I, J) = c_1(I - 2 - 2\log J) + c_2(J-1)^2, \tag{3}$$

where $c_1$ and $c_2$ are the two Lamé parameters.

To characterise the dynamics of the cECM, we follow Onsager's variational formalism [1, 2], which defines the dynamics from a variational principle that satisfies thermodynamic consistency by construction. Towards applying the formalism, we define the Rayleighian

$$\mathcal{R}\left[\boldsymbol{\Phi}; \boldsymbol{V}\right] = \dot{\mathcal{F}}\left[\boldsymbol{\Phi}; \boldsymbol{V}\right] + \mathcal{D}\left[\boldsymbol{\Phi}; \boldsymbol{V}\right], \tag{4}$$

where $\boldsymbol{V} = \partial_t \boldsymbol{\Phi}$ is the three-dimensional velocity of the cECM, $\dot{\mathcal{F}}$ is the rate of change of the free energy, and $\mathcal{D}$ is a dissipation potential characterising the rate of energy dissipation in the system. The rate of change of the free energy can be written as

$$\dot{\mathcal{F}}[\boldsymbol{\Phi}; \boldsymbol{V}] = \int_{\Gamma_R} \left[\frac{1}{2} {S_{\text{el}}}^{ab} \partial_t C_{ab}\right] dS, \tag{5}$$

where ${S_{\text{el}}}^{ab}$ is the elastic component of the second Piola-Kirchhoff stress

$${S_{\text{el}}}^{ab} = 2\frac{\partial w}{\partial C_{ab}}, \tag{6}$$

$\partial_t C_{ab} = \partial_t(\boldsymbol{e}_a \cdot \boldsymbol{e}_b) = \partial_t \partial_a \boldsymbol{\Phi} \cdot \boldsymbol{e}_b + \boldsymbol{e}_a \cdot \partial_t \partial_b \boldsymbol{\Phi} = \partial_a \boldsymbol{V} \cdot \boldsymbol{e}_b + \boldsymbol{e}_a \cdot \partial_b \boldsymbol{V}$, and we have employed Einstein's convention for summation of repeated indices. For a neoHookean material,

$${S_{\text{el}}}^{ab} = 2c_1(G^{ab} - C^{ab}) + 2c_2 J(J-1)C^{ab}, \tag{7}$$

where $G^{ab}$ are the components of the inverse of the metric tensor $G_{ab} = \boldsymbol{e}_{\text{R}a} \cdot \boldsymbol{e}_{\text{R}b}$. We can rewrite Eq. (5) as an integral in the deformed configuration

$$\dot{\mathcal{F}}[\boldsymbol{\Phi}; \boldsymbol{V}] = \int_{\Gamma} {\sigma_{\text{el}}}^{ab} d_{ab} dS, \tag{8}$$

where $\boldsymbol{\sigma}_{\text{el}}$ is the elastic component of the Cauchy stress and $\boldsymbol{d}$ is the rate-of-deformation tensor or strain rate. The components of $\boldsymbol{\sigma}_{\text{el}}$ and $\boldsymbol{d}$ in the convected basis $\{\boldsymbol{e}_1, \boldsymbol{e}_2\}$ coincide with the components of ${S_{\text{el}}}^{ab}/J$ and $\partial_t C_{ab}$ in the basis $\{\boldsymbol{e}_{\text{R}1}, \boldsymbol{e}_{\text{R}2}\}$.

The dissipation potential is given by

$$\mathcal{D}[\boldsymbol{\Phi}; \dot{\boldsymbol{\Phi}}] = \int_{\Gamma} \left\{\eta |\boldsymbol{d}|^2 + \frac{1}{2}\xi \left|\boldsymbol{V} - \boldsymbol{V}_{\text{myo}}\right|^2\right\} dS, \tag{9}$$

where $\eta$ is the bulk viscosity of the cECM, $\xi$ a friction coefficient with the myocardium, $\boldsymbol{V}_{\text{myo}}$ is the velocity of the myocardium, and $|\cdot|$ stands for the 2-norm of a tensor.

The dynamics is then obtained from Onsager's variational principle by minimising the Rayleighian with respect to the velocity

$$\boldsymbol{V} = \arg\min_{\boldsymbol{W}} \mathcal{R}[\boldsymbol{\Phi}; \boldsymbol{W}], \tag{10}$$

subject to constraints. In the case of the cECM, we assume that the movement of the myocardium imposes the normal velocity $\boldsymbol{V} \cdot \boldsymbol{N} = \boldsymbol{V}_{\text{myo}} \cdot \boldsymbol{N}$, where $\boldsymbol{N} = (\boldsymbol{e}_1 \times \boldsymbol{e}_2)/|\boldsymbol{e}_1 \times \boldsymbol{e}_2|$ is the surface normal. This leads to

$$\nabla \cdot \boldsymbol{\sigma} + \boldsymbol{b} = \boldsymbol{0}, \quad (\boldsymbol{V} - \boldsymbol{V}_{\text{myo}}) \cdot \boldsymbol{N} = 0, \tag{11}$$

where $\nabla$ is the covariant derivative on the surface, $\boldsymbol{b} = -\eta \boldsymbol{P}(\boldsymbol{V} - \boldsymbol{V}_{\text{myo}})$ is the frictional force with $P_{\alpha\beta} = \delta_{\alpha\beta} - N_\alpha N_\beta$ the projection operator onto $\Gamma_t$, and $\boldsymbol{\sigma} = \boldsymbol{\sigma}_{\text{el}} + 2\mu \boldsymbol{d}$ is the Cauchy stress tensor. The first equation in Eq. (11) represents force balance tangential to the surface.

### 2 Introducing damage

Eq. (11) characterises a viscoelastic material without damage. In this section, we modify the elastic energy and dissipation potentials of Section 1 to introduce damage. We characterise the amount of damage at a point on $\Gamma_R$ by a scalar field $\alpha(\boldsymbol{X}, t)$ whose value goes from 1 (no damage) to 0 (fully damaged) [4]. The elastic energy Eq. (1) now takes the form

$$\mathcal{F}[\boldsymbol{\Phi}, \alpha] = \int_{\Gamma_R} \left\{ w(I, J) g(\alpha) + \kappa \left[ \omega(\alpha) + \frac{1}{2} l^2 |\nabla \alpha|^2 \right] \right\} dS. \tag{12}$$

Here $g(\alpha)$ is the stiffness damage function, characterising softening of the material due to damage; it should be a convex, monotonically increasing function of $\alpha$ satisfying $g(1) = 1$ and $g(0) = 0$ and with a minimum at $\alpha = 0$. The function $\kappa\omega(\alpha)$ is usually referred to as local dissipated energy density function; $\kappa$ characterises the energy released due to the complete damage of a homogeneous piece of material at the reference configuration (at zero strain). It is a monotonically decreasing function of $\alpha$, with $\omega(1) = 0$ and $\omega(0) = 1$. Finally, $l$ determines a typical length-scale for variations in the damage field. In classical brittle fracture, $\mathcal{G}_c = \kappa / l^2$ measures the critical fracture energy per unit area. Here, following [5], we use

$$g(\alpha) = \frac{\alpha^2}{\alpha^2 + m(1-\alpha)(2-\alpha)}, \qquad \omega(\alpha) = 1 - \alpha, \tag{13}$$

with $m$ measuring the slope of $g(\alpha)$ at $\alpha = 1$, i.e., $g'(1) = m$. $g(\alpha)$ is convex provided $m \geq 3$. The choice for $g(\alpha)$ ensures a smooth transition from undamaged to fully damaged states while maintaining numerical stability. This choice prevents non-physical localization and snap-back instabilities. Moreover, as the internal length scale vanishes, the model converges to a cohesive zone model, where damage localizes into a discrete fracture process zone governed by a traction-separation law. This ensures a physically meaningful transition from diffuse damage to discrete crack formation.

We also modify the dissipation potential

$$\mathcal{D}[\boldsymbol{\Phi}, \alpha; \boldsymbol{V}, \partial_t \alpha] = \int_{\Gamma} \left[ \eta |\boldsymbol{d}|^2 + \frac{1}{2} \xi |\boldsymbol{V} - \boldsymbol{V}_{\text{myo}}|^2 \right] \alpha dS + \int_{\Gamma_R} \lambda (\partial_t \alpha)^2 dS. \tag{14}$$

The first integral is equivalent to Eq. (9), now weighted by $\alpha$ to represent the lower viscosity and friction of a damaged material. The second term represents the rate of energy dissipation due to the rate of change of the damage field characterised by the parameter $\lambda$, with units of viscosity; this parameter is referred to as a kinetic modulus [6]. Although we keep $\lambda$ constant in our simulations, we note that by making $\lambda$ a function of $\boldsymbol{C}$ one could control the different behaviour of crack opening due to stretching instead of compression.

Onsager's variational principle for this model then reads

$$\boldsymbol{V}, \partial_t \alpha = \underset{\boldsymbol{W}, \delta}{\arg\min}\, \mathcal{R}[\boldsymbol{\Phi}, \alpha; \boldsymbol{W}, \delta], \tag{15}$$

where

$$\mathcal{R}[\boldsymbol{\Phi}, \alpha; \boldsymbol{V}, \partial_t \alpha] = \dot{\mathcal{F}}[\boldsymbol{\Phi}, \alpha; \boldsymbol{V}, \partial_t \alpha] + \mathcal{D}[\boldsymbol{\Phi}, \alpha; \boldsymbol{V}, \partial_t \alpha], \tag{16}$$

subject to the constraints

$$\boldsymbol{V} \cdot \boldsymbol{N} = v_n, \qquad \partial_t \alpha > 0. \tag{17}$$

The rate of change of the free energy is now

$$\dot{\mathcal{F}}[\boldsymbol{\Phi}, \alpha; \boldsymbol{V}, \partial_t \alpha] = \int_{\Gamma} \left[ g(\alpha) {\sigma_{\text{el}}}^{ab} d_{ab} \right] dS + \int_{\Gamma_R} \left[ w g'(\alpha) + \kappa \left( \omega'(\alpha) - l^2 \Delta \alpha \right) \right] dS. \tag{18}$$

Solving for Eq. (15), we recover Eq. (11), now with $\boldsymbol{\sigma} = g(d) \boldsymbol{\sigma}_{\text{el}} + 2\mu\alpha \boldsymbol{d}$ and $\boldsymbol{b} = -\eta\alpha \boldsymbol{P}(\boldsymbol{V} - \boldsymbol{V}_{\text{myo}})$. We also get the evolution equation for the phase field

$$\partial_t \alpha = \min \left( 0, -\frac{1}{\lambda} \left[ w g'(\alpha) + \kappa \left( \omega'(\alpha) - l^2 \Delta \alpha \right) \right] \right). \tag{19}$$

The critical energy density is then given by

$$w_{\text{crit}} = \frac{\kappa}{m}. \tag{20}$$

| Name | Symbol | Units | Non-dimensional equivalent | Range in simulations |
|---|---|---|---|---|
| Lamé parameters | $c_1, c_2$ | Pa m | $1, \tilde{c}_2 = \frac{c_2}{c_1}$ | $1, 1$ |
| Critical energy density | $w_{\text{crit}}$ | Pa m | $\tilde{w}_{\text{crit}} = \frac{w_{\text{crit}}}{c_1}$ | 0.16 |
| Elastic energy release | $\kappa$ | Pa m | $\tilde{\kappa} = \frac{\kappa}{c_1}$ | 0.5 |
| Fracture sharpness | $l$ | m | $\tilde{l} = \frac{l}{R}$ | $4.47 \cdot 10^{-2}$ |
| Viscosity | $\mu$ | Pa m s | $\tilde{\mu} = \frac{\mu}{\lambda}$ | $10^{-4}$ |
| Friction | $\eta$ | Pa m$^{-1}s$ | $\tilde{\eta} = \frac{\eta R^2}{\lambda}$ | 50 |
| Damage dissipation | $\lambda$ | Pa m s | 1 | 1 |
| Typical domain radius | $R$ | m | 1 | 1 |
| Beating frequency | $\omega$ | s$^{-1}$ | $\tilde{\omega} = \frac{\lambda}{c_1}\omega$ | 2.22 |
| Penalty reheal | $K_p$ | Pa m | $\tilde{K}_p = K_p/c_1$ | $2 \cdot 10^3$ |
| Penalty shape | $K_s$ | Pa m | $\tilde{K}_s = K_p/c_1$ | $10^5$ |

Table 1: Model parameters, non-dimensionalisation, and values used in simulations.

### 3 Dimensional analysis

The features of the fracture patterns that emerge from the model in Section 2 depend on the different parameters of the model. In Table 1 we summarise the different parameters and non-dimensionalise the system by taking units of tension relative to $c_1$, units of time relative to $\lambda/c_1$, and units of length relative to the typical domain size $R$. We also specify the parameters used for all simulations in the manuscript.

### 4 Boundary conditions

In simulations, we analyze the cECM within a region bounded by the reconstructed shape of the myocardium, obtained through nuclear tracking. This process results in surfaces with two open ends, resembling a cylinder. At both ends, we impose Dirichlet boundary conditions for the myocardium's velocity and homogeneous Neumann boundary conditions for the damage field,

$$\boldsymbol{V} = \boldsymbol{V}_{\text{myo}}, \qquad \nabla\alpha = 0, \qquad \text{at } \partial\Gamma_t. \tag{21}$$

### 5 Numerical implementation

#### 5.1 Time discretisation

We first describe the time discretisation of the equations. We discretise time in a non-uniform grid $[t^{(0)}, t^{(1)}, \ldots, t^{(N)}]$ with time steps $\Delta t^{(N)} = t^{(N+1)} - t^{(N)}$. We then define $\boldsymbol{\Phi}^{(n)}(\boldsymbol{X})$ and $\alpha^{(n)}(\boldsymbol{X})$ as the deformation and damage fields at the time-step $n$. Given those values, we can find the $\boldsymbol{\Phi}^{(n+1)}(\boldsymbol{X})$ and $\alpha^{(n+1)}(\boldsymbol{X})$ by solving a dicrete Onsager variational principle

$$\boldsymbol{\Phi}^{(n+1)}, \alpha^{(n+1)} = \arg\min_{\boldsymbol{W},\delta} \mathcal{R}[\boldsymbol{\Phi}^{(n)}, \alpha^{(n)}; \boldsymbol{W}, \delta]. \tag{22}$$

with Rayleighian

$$\begin{aligned}\mathcal{R}[\boldsymbol{\Phi}^{(n)}, \alpha^{(n)}; \boldsymbol{\Phi}^{(n+1)}, \alpha^{(n+1)}] =& \frac{1}{\Delta t^{(N)}} \left(\mathcal{F}[\boldsymbol{\Phi}^{(n+1)}, \alpha^{(n+1)}] - \mathcal{F}[\boldsymbol{\Phi}^{(n)}, \alpha^{(n)}]\right) \\ &+ \mathcal{D}\left[\boldsymbol{\Phi}^{(n)}, \alpha^{(n)}; \frac{\boldsymbol{\Phi}^{(n+1)} - \boldsymbol{\Phi}^{(n)}}{\Delta t^{(n)}}, \frac{\alpha^{(n+1)} - \alpha^{(n)}}{\Delta t^{(n)}}\right].\end{aligned} \tag{23}$$

Note that in the discretisation of the time-derivative of the free energy, the second term (free energy in the previous time-step) does not depend on the target variables $\boldsymbol{\Phi}^{(n+1)}, \alpha^{(n+1)}$ and can thus be discarded.

#### 5.2 Weakly imposition of constraints through penalty functions

To deal with the constraints in Eq. (17), we add two extra free energies at step $(n+1)$. We first add an energy to ensure that the shape of the cECM follows that of the myocardium (which is given as data)

$$\mathcal{F}_{\text{shape}}(\boldsymbol{\Phi}^{(n)}; \boldsymbol{\Phi}^{(n+1)}) = \int_{\Gamma^{(n)}} K_s \left[\left(\boldsymbol{\Phi}^{(n+1)} - \boldsymbol{\Xi}^{(n+1)}\left(\boldsymbol{\Phi}^{(n)}\right)\right) \cdot \boldsymbol{N}^{(n)}\right]^2 dS, \tag{24}$$

where $\boldsymbol{\Xi}^{(n)}(\boldsymbol{x})$ is the closest-point projection of $\boldsymbol{x}$ onto the shape of the myocardium at time-step $(n)$. Thus this function penalises the motion of $\boldsymbol{\Phi}^{(n+1)}$ relative to the closest point projection of $\boldsymbol{\Phi}^{(n)}$ onto the myocardium along the normal to the surface.

We also add an energy to ensure irreversibility of the damage process as follows

$$\mathcal{F}_{\text{irrever}}(\alpha^{(n)}; \alpha^{(n+1)}) = \int_{\Gamma_R} \Theta\left(\alpha^{(n+1)}; \alpha_{\text{min}}^{(n)}\right) dS, \tag{25}$$

where

$$\Theta(\alpha; \alpha_{\text{min}}) = \left\{ \begin{array}{ll} \frac{1}{3} K_p (\alpha - \alpha_{\text{min}})^3 & \text{if } \alpha < \alpha_{\text{min}}, \\ 0 & \text{otherwise.} \end{array} \right., \tag{26}$$

and $\alpha_{\text{min}}^{(n)}(\boldsymbol{X}) = \min\{\alpha^{(1)}(\boldsymbol{X}), \ldots, \alpha^{(n)}(\boldsymbol{X})\}$. This function penalises values of $\alpha < \alpha_{\text{min}}$ weakly through the penalty parameter $K_p$; in the limit of $K_p \to \infty$, this ensures that irreversibility of damage is satisfied.

#### 5.3 Spatial discretisation

To discretise in space, we follow a usual finite element discretisation. The surface is described by a triangular mesh and the fields driscretised as

$$\boldsymbol{\Phi}^{(n)}(s^1, s^2) = \sum_I \boldsymbol{x}_I^{(n)} B_I(s^1, s^2), \qquad \alpha^{(n)}(s^1, s^2) = \sum_I \alpha_I^{(n)} B_I(s^1, s^2), \tag{27}$$

where $\boldsymbol{x}_I^{(n)}$ and $\alpha_I^{(n)}$ are the position and the damage at node $I$ at time-step $n$, and $B_I$ are the basis functions, which are the usual linear basis functions of first-order Lagrangian interpolation. Eq. (22) can then be written as

$$\left\{\boldsymbol{\Phi}_I^{(n+1)}, \alpha_I^{(n+1)}\right\}_{I=1}^{N} = \underset{\boldsymbol{W}, \delta}{\arg\min}\, \mathcal{R}[\boldsymbol{\Phi}^{(n)}, \alpha^{(n)}; \boldsymbol{W}, \delta], \text{ where } \boldsymbol{W} = \sum_I \boldsymbol{W}_I^{(n)} B_I,\ \delta = \sum_I \delta_I^{(n)} B_I. \tag{28}$$

The integrals in the expressions of the free energy, and dissipation potential are calculated by summing contributions from each one of the triangles of the mesh and using Gaussian integration with 3 integration points in each of the triangles. We solve this minimisation process using hiperlife [7]. The code can be found here https://git.embl.de/grp-torres-sanchez/cECM-fractures.

#### References


[1] Masao Doi. “Onsager’s variational principle in soft matter”. en. In: *J. Phys. Condens. Matter* 23.28 (20 7 2011), p. 284118.

[2] Marino Arroyo et al. “Onsager’s Variational Principle in Soft Matter: Introduction and Application to the Dynamics of Adsorption of Proteins onto Fluid Membranes”. en. In: *The Role of Mechanics in the Study of Lipid Bilayers*. Ed. by David J Steigmann. CISM International Centre for Mechanical Sciences. Cham: Springer International Publishing, 2018, pp. 287–332.

[3] Jerrold E Marsden and Thomas J R Hughes. *Mathematical Foundations of Elasticity*. en. New York, New York, USA: Courier Corporation, 1994.

[4] Jean-Jacques Marigo, Corrado Maurini, and Kim Pham. “An overview of the modelling of fracture by gradient damage models”. In: *Meccanica* 51.12 (Jan. 2016), pp. 3107–3128.

[5] Eric Lorentz, S Cuvilliez, and K Kazymyrenko. “Convergence of a gradient damage model toward a cohesive zone model”. en. In: *Comptes Rendus Mécanique* 339.1 (Dec. 2010), pp. 20–26.

[6] Rudy J M Geelen et al. “A phase-field formulation for dynamic cohesive fracture”. In: *Comput. Methods Appl. Mech. Eng.* 348 (Jan. 2019), pp. 680–711.

[7] Daniel Santos-Oliván, Guillermo Vilanova, and Alejandro Torres-Sánchez. *hiperlife*. Version release-5.0.0-alpha. Feb. 2025. URL: http://dx.doi.org/10.5281/zenodo.14927572.
